# A comparison of single-cell trajectory inference methods: towards more accurate and robust tools

**DOI:** 10.1101/276907

**Authors:** Wouter Saelens, Robrecht Cannoodt, Helena Todorov, Yvan Saeys

**Affiliations:** Data mining and Modelling for Biomedicine, VIB Center for Inflammation Research, Ghent, Belgium.; Department of Applied Mathematics, Computer Science and Statistics, Ghent University, Ghent, Belgium.; Center for Medical Genetics, Ghent University Hospital, Ghent, Belgium.; Centre International de Recherche en Infectiologie, Inserm, U1111, Université Claude Bernard Lyon 1, CNRS, UMR5308, École Normale Supérieure de Lyon, Univ Lyon, F-69007, Lyon, France

## Abstract

Using single-cell-omics data, it is now possible to computationally order cells along trajectories, allowing the unbiased study of cellular dynamic processes. Since 2014, more than 50 trajectory inference methods have been developed, each with its own set of methodological characteristics. As a result, choosing a method to infer trajectories is often challenging, since a comprehensive assessment of the performance and robustness of each method is still lacking. In order to facilitate the comparison of the results of these methods to each other and to a gold standard, we developed a global framework to benchmark trajectory inference tools. Using this framework, we compared the trajectories from a total of 29 trajectory inference methods, on a large collection of real and synthetic datasets. We evaluate methods using several metrics, including accuracy of the inferred ordering, correctness of the network topology, code quality and user friendliness. We found that some methods, including Slingshot, TSCAN and Monocle DDRTree, clearly outperform other methods, although their performance depended on the type of trajectory present in the data. Based on our benchmarking results, we therefore developed a set of guidelines for method users. However, our analysis also indicated that there is still a lot of room for improvement, especially for methods detecting complex trajectory topologies. Our evaluation pipeline can therefore be used to spearhead the development of new scalable and more accurate methods, and is available at github.com/dynverse/dynverse.

To our knowledge, this is the first comprehensive assessment of trajectory inference methods. For now, we exclusively evaluated the methods on their default parameters, but plan to add a detailed parameter tuning procedure in the future. We gladly welcome any discussion and feedback on key decisions made as part of this study, including the metrics used in the benchmark, the quality control checklist, and the implementation of the method wrappers. These discussions can be held at github.com/dynverse/dynverse/issues.

## Introduction

Single-cell -omics technologies now make it possible to model biological systems more accurately than ever before^1^. One area where single-cell data has been particularly useful is in the study of cellular dynamic processes, such as the cell cycle, cell differentiation and cell activation^2^. Such dynamic processes can be computationally modelled using trajectory inference (TI) methods (also known as pseudemporal ordering methods), which use single-cell profiles from a population in which the cells are at different unknown points in the dynamic process^3,4,5^. These methods computationally order the cells along a trajectory topology, which can be linear, bifurcating, or a more complex tree or graph structure. Because TI methods offer an unbiased and transcriptome-wide understanding of a dynamic process^1^, they allow the objective identification of new (primed) subsets of cells^6^, delineation of a differentiation tree^7,8^ and inference of regulatory interaction responsible for one or more bifurcations^9^. Current applications of TI focus on specific subsets of cells, but ongoing efforts to construct transcriptomic catalogues of whole organisms^10,11^ underline the urgency for accurate, scalable^9,12^ and user-friendly TI methods.

**Table 1.**
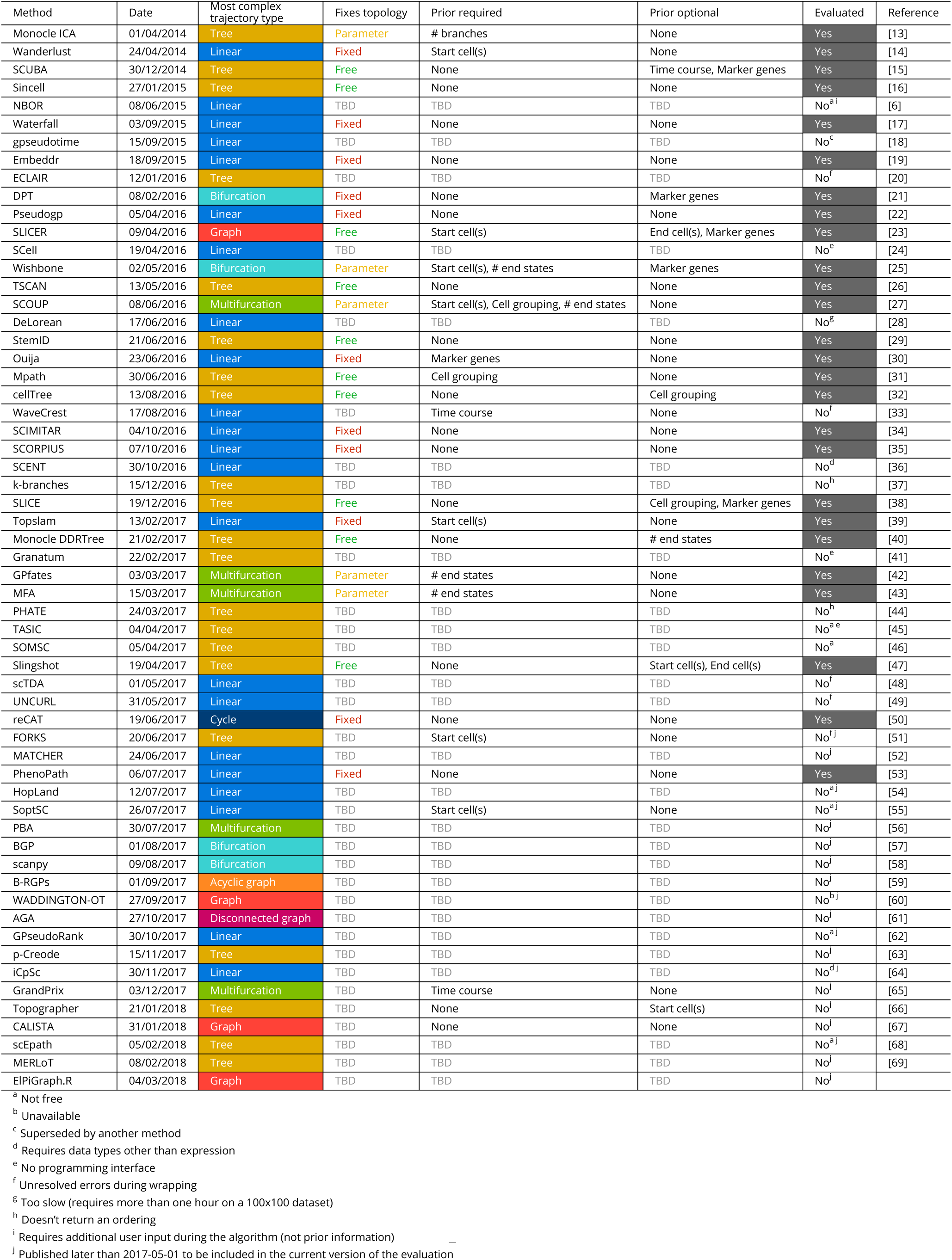
Overview of the trajectory inference methods included in this study, and several characteristics thereof. This table will be continuously updated online.

A plethora of TI methods has been developed over the last years, and even more are being created every month (**Supplementary Figure 1a**). It is perhaps surprising that of the 59 methods in existence today, almost all methods have a unique combination of characteristics (**Table 1**), in terms of the required inputs (prior information), produced outputs (topology fixing and trajectory type) and methodology used (not shown). One distinctive characteristic of TI methods is whether the topology of the trajectory is inferred computationally, or was fixed by design. Early TI methods typically fixed the topology algorithmically (e.g. linear^14,6,17,18^ or bifurcating^21,25^), or through parameters provided by the user^13,27^. These methods therefore mainly focused on correctly ordering the cells along this fixed topology. Other methods attempt to infer the topology computationally, which increases the difficulty of the problem at hand, but allows these methods to be broadly applicable on more use cases. Methods that perform topology inference are still in the minority, though current trends suggest this will soon change (**Supplementary Figure 1c**). A particularly interesting development is presented in the AGA method^61^ which is the only TI method currently able to deal with disconnected graphs.

**Figure 1.**
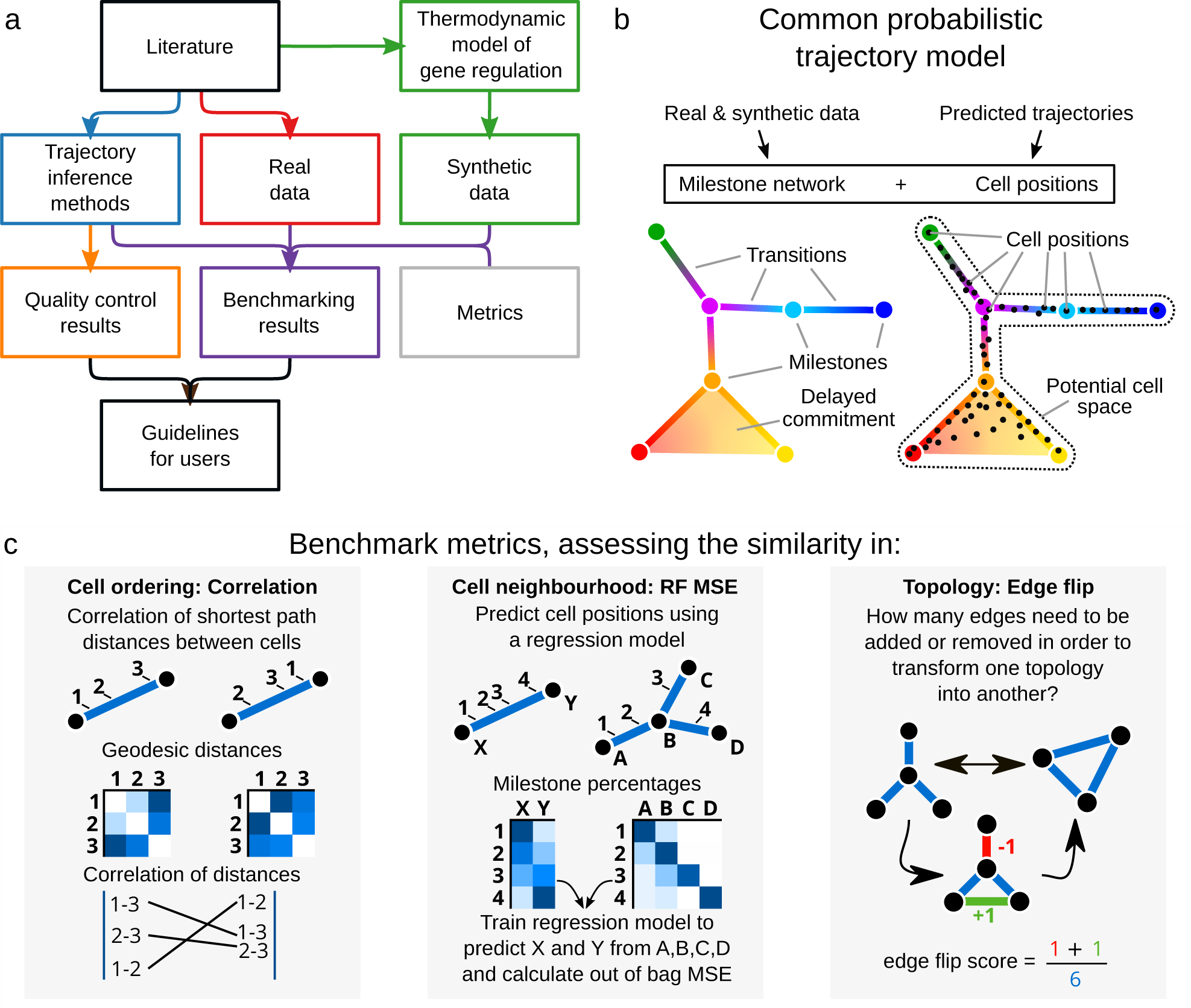
An overview of several key aspects of the evaluation. a) A schematic overview of our evaluation pipeline. b) In order to make any two trajectories comparable with one another, a common trajectory model was used to represent gold standard trajectories from the real and synthetic datasets, as well as any predicted trajectories from TI methods. c) Three metrics were defined in order to assess the similarity in cell ordering, cell neighbourhood, and topology, for any two trajectories.

Another key characteristic of TI methods is the selection of prior information that a method requires or can optionally exploit. Prior information can be supplied as a starting cell from which the trajectory will originate, a set of important marker genes, or even a grouping of cells into cell states. Providing prior information to a TI method can be both a blessing and a curse. In one way, prior information can help the method to find the correct trajectory among many, equally likely, alternatives. On the other hand, incorrect or noisy prior information can bias the trajectory towards current knowledge. Moreover, prior information is not always easily available, and its subjectivity can therefore lead to multiple equally plausible solutions, restricting the applicability of such TI methods to well studied systems.

A reductionist approach to characterising TI methods consists in dissecting them into a set of algorithmic components, as any component can have a significant impact on the performance, scalability, and output data structures. Across all TI methods, these components can be broadly grouped into two stages; (i) conversion to a simplified representation using dimensionality reduction, clustering or graph building and (ii) ordering the cells along the simplified representation^4^. Interestingly, components are frequently shared between different algorithms (**Supplementary Figure 2**). For example, minimal spanning trees (MST), used in the first single-cell RNA-seq trajectory inference methods^13^, is shared by almost half of the methods we evaluated (**Supplementary Figure 2b**).

Given the diversity in TI methods, an important issue to address is a quantitative assessment of the performance and robustness of the existing TI methods. Many attempts at tackling this issue have already been made^21,26,23,27,32,35,42,43^, but due to the high number of TI methods available today and the great diversity in the outputted data structures, a comprehensive benchmarking evaluation of TI methods is still lacking. This is problematic, as new users to the field are confronted with a wide array of TI methods, without a clear idea about what method would be the most optimal for their problem. Moreover, the strengths and weaknesses of existing methods need to be assessed, so that new developments in the field can focus on improving the current state-of-the-art.

## Results

In this study, we performed a comprehensive evaluation for 29 TI methods (Overview: **Figure 1a**, Extended overview: **Supplementary Figure 3**). The inclusion criterion for TI methods was primarily based on their free availability, the presence of a programming interface, and the date of publication (**Table 1**). Only methods published before june 2017 are included in the current version of the evaluation, while more recent methods will be added in the next version. The evaluation comprised three core aspects: (i) source-code and literature-based characterisation of TI methods, (ii) assessment of the accuracy and scalability of TI methods by comparing predicted trajectories with a gold standard, and (iii) a quality control of the provided software and documentation.

In order to make gold standard trajectories and predicted trajectories directly comparable to one another, we developed a common probabilistic model for representing trajectories from all possible sources (**Figure 1b**). In this model, the overall topology is represented by a network of “milestones”, and the cells are placed within the space formed by each set of connected milestones. We defined a set of metrics for comparing the likeness of such trajectories, each assessing a different aspect of the trajectory: the similarity in cell ordering, the cell neighbourhood and the topology (**Figure 1c**). The data compendium consisted of both synthetic datasets, which offer the most exact gold standard, and real datasets, which offer the highest biological relevance. These real datasets came from a variety of single-cell technologies, organisms, and dynamic processes, and contain several types of trajectory topologies (**Supplementary Figure 4** and **Supplementary Table 1**). To generate synthetic datasets, we simulated a gene regulatory network using a thermodynamic model of gene regulation^70^, and subsequently simulated a single-cell profiling experiment by matching the distributions of the synthetic data with real reference datasets (**Supplementary** Figure 5).

As the aim of this study is to provide both guidelines for the use of TI methods and a base for the development of new TI methods, we not only assessed the accuracy of the predictions made by a trajectory, but also investigated the quality of the method’s implementation. To do this, we scored each method using a checklist of important scientific and software development practices, including software packaging, documentation, automated code testing, and peer review. This allowed us to rank each method based on its user friendliness, developer friendliness, and potential broad applicability on new unseen datasets. Finally, using both the benchmark results and quality control, we produced a flow chart with practical guidelines for selecting the most appropriate TI method for a given use case.

### Evaluation of trajectory inference methods

An overview of the main results from this study is shown in **Figure 2**. This includes an overview of the results obtained from the method characterisation (**Figure 2a**), the benchmarking evaluation (**Figure 2b**), and the quality control evaluation (**Figure** 2c).

**Figure 2.**
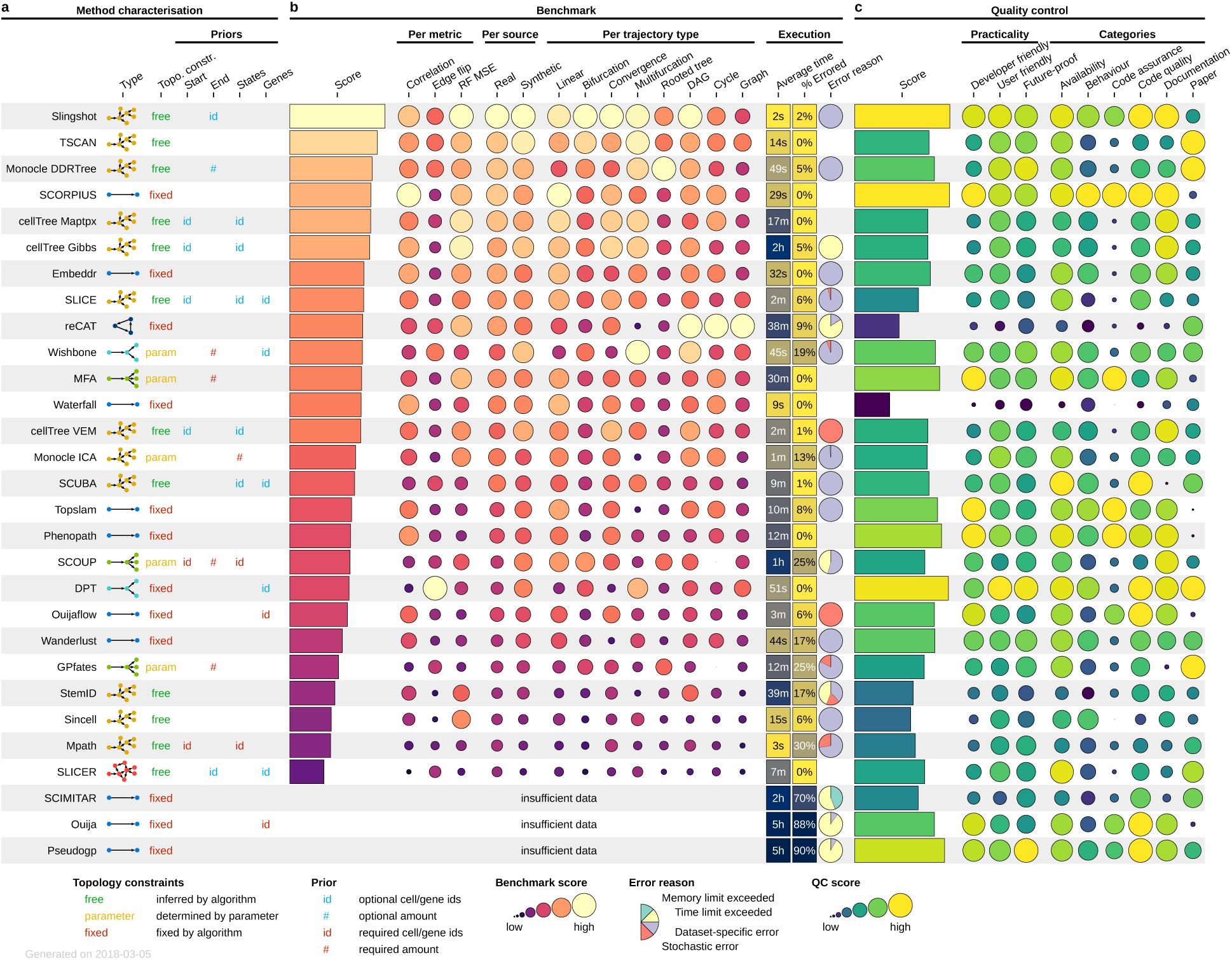
Overview of the results on the evaluation of 29 TI methods. a) The methods were characterised according to the most complex trajectory type they can infer, whether or not the inferred topology is constrained by the algorithm or a parameter, and which prior information a method requires or can optionally use. b) The methods are ordered according to the overall score achieved in the benchmark evaluation. Also shown are the aggregated scores per metric, source and trajectory type, as well as the average execution time across all datasets and the percentage of executions in which no output was produced. c) Overall performance in the quality control evaluation is highly variable, even amongst the highest ranked methods according to the benchmark evaluation. Also listed are the quality control scores aggregated according to practicality and the different categories.

Having ordered all methods by their overall benchmarking score, we found that Slingshot predicted the most accurate trajectories, followed by TSCAN and Monocle DDRTree. When we looked at the benchmark scores per trajectory type, Slingshot was the only method that performed well across most trajectory types. However, we found that several methods were specialised in predicting specific trajectory types; for example SCORPIUS for linear trajectories, reCAT for cycles, and Monocle DDRTree for trees.

We observed a high correlation (0.7-0.9) between results originating from real datasets versus those originating from synthetic datasets (**Supplementary Figure 6**). This confirms both the relevance of the synthetic data and the accuracy of the gold stan-dard in the real datasets. However, this correlation was lower for converging and multifurcating datasets, potentially because only a small number of real datasets were available for such topologies (**Supplementary Table 1**).

As the different metrics were selected to assess the correctness of a trajectory in various approaches, it is expected to observe differences in rankings of TI methods amongst the different metrics. While we saw no direct link between the edge flip scores versus the correlation or RF MSE metric, methods that obtained a high correlation score also tended to obtain higher RF MSE scores (**Supplementary Figure 7**). In addition, the methods that were able to detect more complex trajectory types also obtained higher edge flip scores, in comparison to methods whose trajectory topology was fixed to a simple trajectory topology. We will explore this issue in more detail in a further section.

During the execution of a method on a dataset, if the time limit (>6h) or memory limit (32GB) was exceeded, or an error was produced, a zero score was returned for that execution. If a method consistently generated errors on this dataset across all replicates, the error is called “data-specific”, otherwise “stochastic”. Several methods obtained a high overall score despite having 5 to 10% of failed executions (e.g. Monocle DDRTree, SLICE, Wishbone), meaning these methods could rank even higher if not for the failed executions. Methods that do not scale well with respect to the number of cells or genes will exceed the time or memory limits for the largest datasets (reCAT, SCOUP, StemID). For a few methods, the time or memory limits were exceeded too often, making the benchmarking results uncomparable to those of other methods (SCIMITAR, Ouija, Pseudogp).

### Trajectory types and topology complexity

In most cases, the methods which were specifically designed to handle a particular trajectory type, also performed better on data containing this particular trajectory type (**Supplementary Figure 8**). These methods had typically better edge flip scores - as can be expected - and RF MSE scores, compared to methods not able to handle the particular trajectory type. However, the correlation score typically followed the opposite pattern, where methods restricted to linear trajectory types, such as embeddr, SCORPIUS and phenopath, produced the best ordering, irrespective of whether the dataset contained a linear trajectory or more complex trajectories. To further investigate the effect of trajectory complexity on performance, we divided them in two groups: linear methods (restricted to linear and cyclic topologies) and non-linear methods. While there were no significant differences in performance on linear datasets, non-linear methods had significantly higher edge flip scores but lower correlation scores on datasets containing more complex trajectory types (**Supplementary Figure 9**). Together, this indicates that current non-linear methods potentially contain less accurate ordering algorithms. A combination of the ordering methods from the top linear methods, combined with the topology inference from top non-linear methods, could therefore be a possibility for future research.

Despite their similar overall performance, the topologies predicted by the top methods differed considerably. Trajectories detected using the default parameters of slingshot and cellTree tended to be simpler, while those detected by TSCAN and Monocle DDRTree gravitated towards more complex topologies (**Figure 3**). Monocle DDRTree, for example frequently predicted a tree topology, even when only a cyclic, linear or bifurcating topology was present in the data (**Figure 3a**). Trajectories generated by Monocle DDRTree (at default parameters) tended to contain more nodes and edges (**Figure 3b**), which could give this method an advantage on datasets with complex tree topologies, but could also explain its relatively low performance on linear and bifurcating datasets. Indeed, when we assessed how often a method can infer the correct topology, slingshot and TSCAN were good at correctly predicting linear, bifurcating and converging trajectories, while Monocle DDRTree was by far the best method to infer tree topologies. Nonetheless, the overall accuracy of topology prediction was very low, with Slingshot correctly predicting bifurcating and converging topologies for half of the datasets, and Monocle DDRTree predicting the correct tree topology in 12% of the cases (**Supplementary Figure 10**). Inferring the correct topology without parameter tuning, is therefore still an open challenge. Conversely, when the data contains a complex trajectory structure, TI will currently still require a considerable guidance by the end user, to either optimise the parameters or to choose the method with which the output best fits the user’s

**Figure 3.**
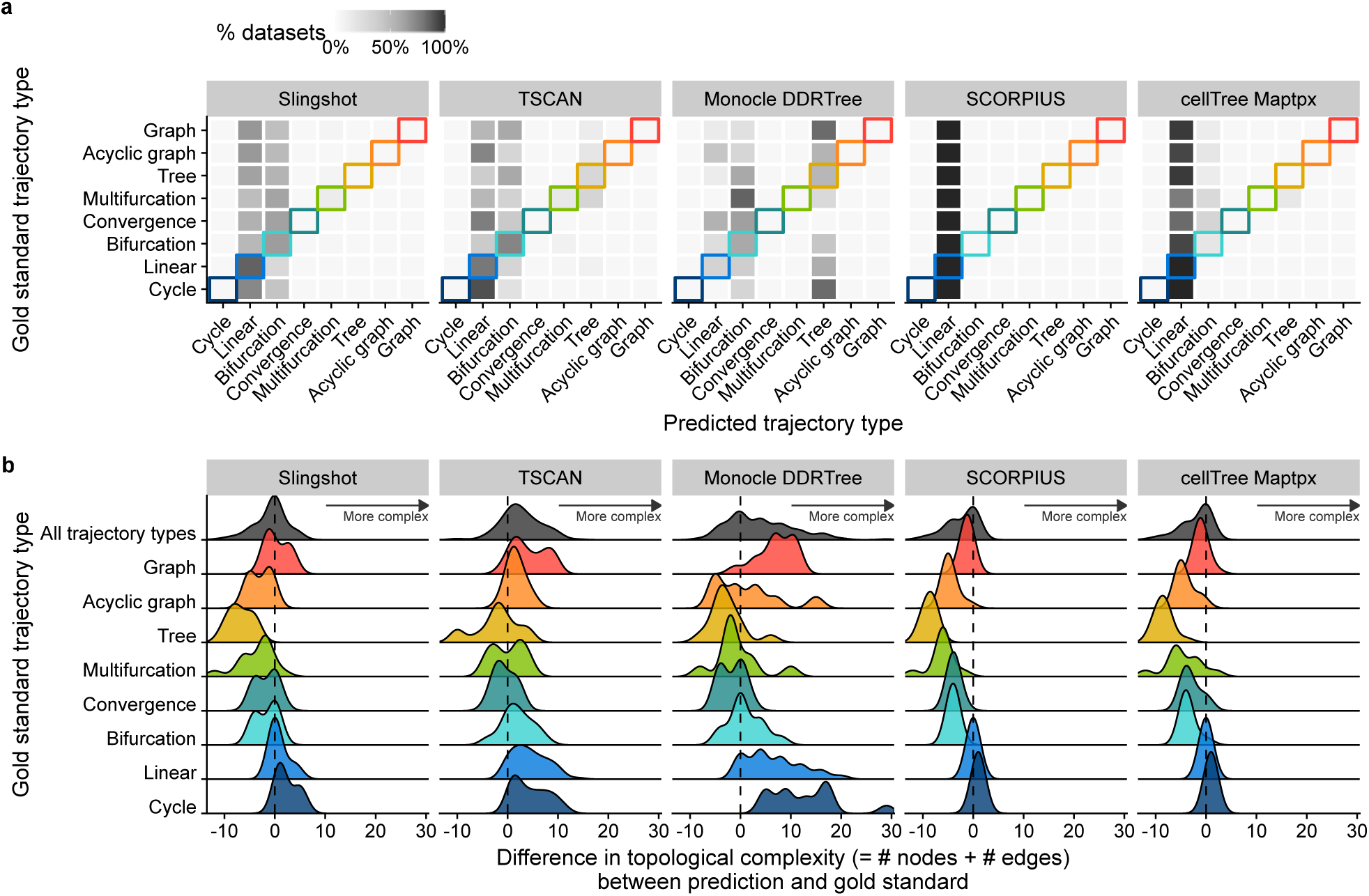
Comparing the ability of the top TI methods to detect the correct trajectory type. a) % of datasets on which a TI method detects a particular trajectory type, compared with the correct trajectory type. b) Distributions of the topological complexity, defined by the sum of the number of milestones and edges between milestones, compared with the true trajectory type present in the data.

### Effect of prior information

In the current version of the evaluation, we only provided prior information when a method required it. We did not observe a major difference in method performance between methods which did and did not receive prior information (**Supplementary Figure 11**). Rather, methods which received prior information were on average positioned in the middle of the ranking. Furthermore, we could not find any dataset where methods which received prior information performed significantly better than other methods.

### Algorithm components

We assessed whether the components of an algorithm could be predictive of a method’s performance, using both random forest classification and statistical testing. Methods which included principal curves (such as Slingshot and SCORPIUS), k-means (which include Slingshot and several other top scoring methods) and some graph building (which include almost every top scoring method) tended to have a slightly higher performance (**Supplementary Figure 12** and **Supplementary Table 2**). On the other hand, methods using t-SNE and ICA for their dimensionality reduction were ranked lower, although this was not statistically significant.

### Method quality control

While not directly related to the accuracy of the inferred trajectory, the quality of the implementation is also an important evaluation metric. Good unit testing assures that the implementation is correct, good documentation makes it easier for potential users to apply the method on their data, and overall good code quality makes it possible for other developers to adapt the method and extend it further. We therefore looked at the implementation of each method, and assessed its quality using a transparent scoring scheme (**Supplementary Table 3**). The individual quality checks can be grouped in two ways: what aspect of the method they investigate (availability, code quality, code assurance, documentation, method’s behaviour at runtime and the depth by which the method was presented in its study) or which purpose(s) they serve (user friendliness, developer friendliness or future proof). These categorisations can help current developers to improve their tool, and guide the selection of users and developers to use these tools for their purpose. After publishing this preprint, we will contact the authors of each method, allowing them to improve their method before the final publishing of the evaluation.

We found that most methods fulfilled the basic criteria, such as free availability and basic code quality criteria (**Figure 4**). We found that recent methods had a slightly better quality than older methods (**Supplementary Figure 13**). However, several quality aspects were consistently lacking for the majority of the methods (**Figure 4 right**) and we believe that these should receive extra attention from developers. Although these outstanding issues cover all five categories, code assurance and documentation in particular are problematic areas, notwithstanding several studies pinpointing these as good practices^71,72^.

**Figure 4.**
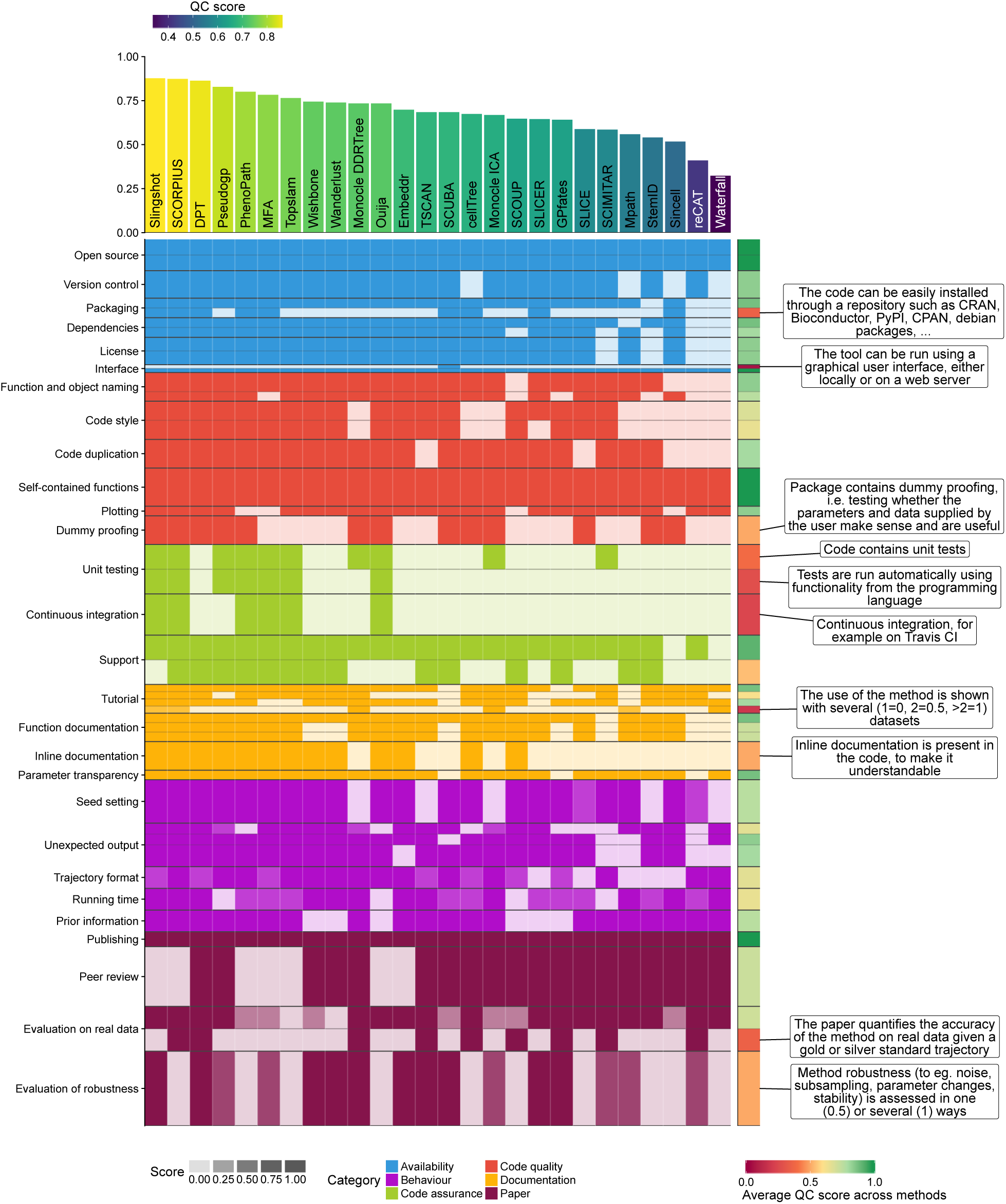
Overview of the quality control results for every method. Shown is the score given for each methods on every item from our quality control score sheet (**Supplementary Table 3**). Each aspect of the quality control was part of a category, and each category was weighted so that it contributed equally to the final quality score. Within each category, each aspect also received a weight depending on how often it was mentioned in a set of papers discussing good practices in tool development and evaluation. This weight is represented in the plot as distance on the y-axis. Top: Average QC score for each method. Right: The average score of each quality control item. Shown into more detail are those items which had a score lower than 0.5.

We observed no clear relation between method quality and method performance (**Figure 2** and **Supplementary Figure 14**). We could also not find any quality aspect which was significantly associated with method performance.

### Practical guidelines

Based on the results of our benchmark, we created a set of practical guidelines for method users **Figure 5**. As a method’s performance is heavily dependent on the trajectory type being studied, the choice of method will by primarily driven by the prior knowledge of the user about what trajectory topology is expected in the data. For the majority of use cases, the user will know very little about the expected trajectory, except perhaps whether the data is expected to contain multiple trajectories, cycles or a complex tree structure. In each of these use cases, a different set of methods performed optimally, with Monocle DDRTree performing best when the data contained a complex tree, and Slingshot performing equally well on less complex trajectories No methods dealing with multiple trajectories or cycles were included in the current version of the evaluation, although AGA^61^ for disconnected trajectories, and Waddington-OT^60^ and AGA^61^ for regular graphs are currently the only methods in literature able to handle these types of trajectories. In the case where the user would know the exact expected topology, our evaluation suggests the use of reCAT for cycles, SCORPIUS for linear trajectories, and Slingshot for bifurcating trajectories, although Slingshot could return other topologies if it would fit the data more accurately. The most difficult use case is when the topology is known but more complex than a bifurcating or cyclic trajectory. Here, to our knowledge, only ElPiGraph.R github.com/Albluca/ElPiGraph.R, which is not yet published, can be used.

**Figure 5.**
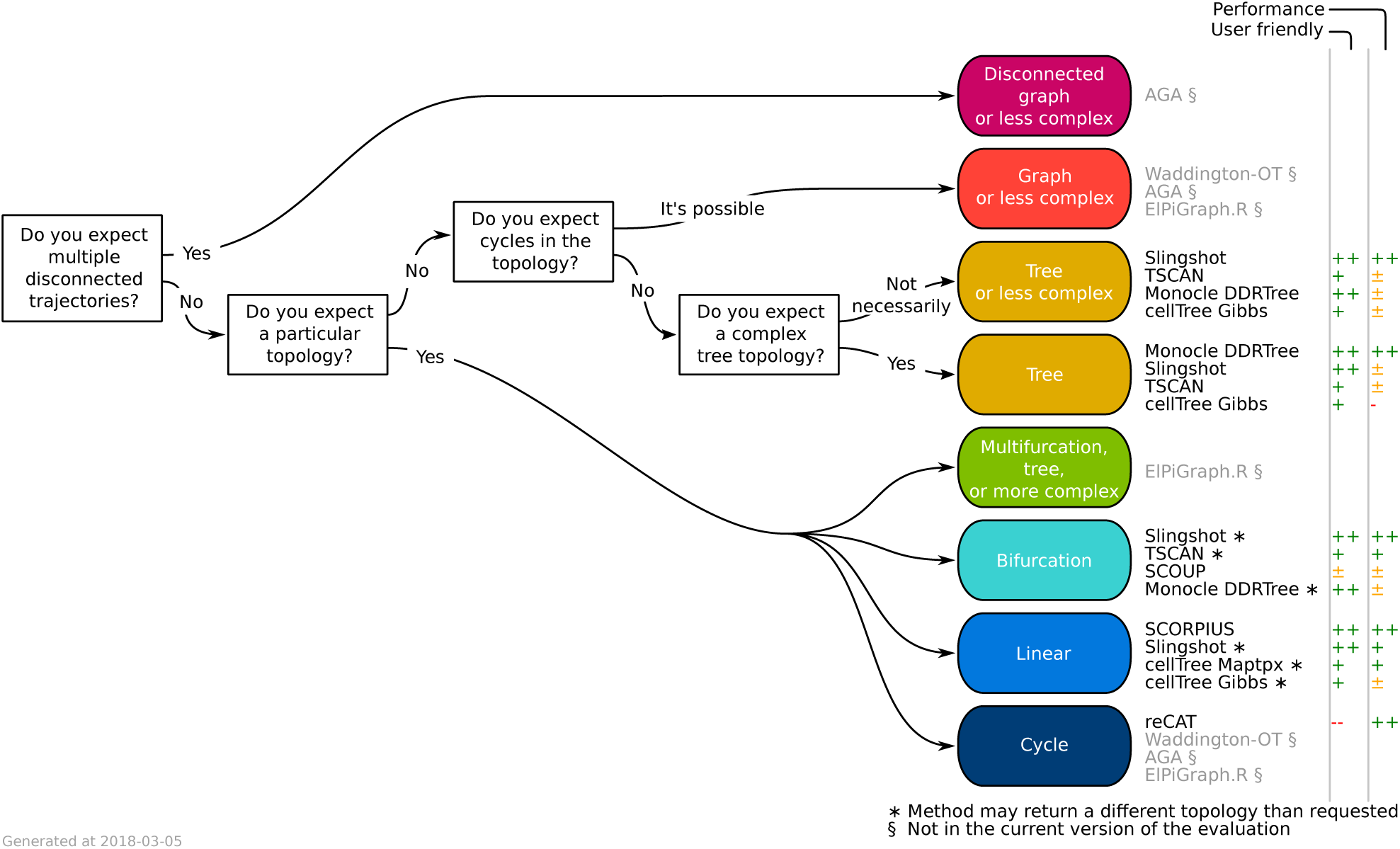
Practical guidelines for method users. As the performance of a method most heavily depended on the topology of the trajectory, the choice of TI method will be primarily influenced by the user’s existing knowledge about the expected topology in the data. We therefore devised a set of practical guidelines, which combines the method’s performance, user friendliness and the number of assumptions a user is willing to make about the topology of the trajectory. Methods to the right are ranked according to overall performance. Further to the right are shown the user friendly scores (++: ≥ 0.9, +: ≥ 0.8, ± ≥ 0.65, - ≥ 0.5) and overall performance (++: top method, +: difference between top method’s performance ≥ −0.05, ±: ≥-0.2, -: ≥ −0.5).

When choosing a method, it is important to take two further points into account. First, it is important that a trajectory and the downstream results and/or hypotheses originating from it are confirmed by multiple TI methods. This to make sure the model is not biased towards the particular model underlying a TI method, for example its preferred trajectory type. Second, even if the expected topology is known, it can be beneficial to also try out methods which make the less assumptions about the trajectory topology. When the expected topology is confirmed using such a method, it provides extra evidence to the user’s knowledge of the data. When a more complex topology is produced, this could indicate the presence of a more complex trajectory in the data than was expected by the user.

## Discussion

Trajectory inference is unique among most other categories of single-cell analysis methods, such as clustering, normalisation and differential expression, because it models the data in a way that was almost impossible using bulk data. Indeed, for no other single-cell analysis types have so many tools been developed, according to several repositories such as omictools.org^73^, the “awesome single cell software” list^74^ and scRNA-tools.org^75^. It is therefore critical that these methods, now reaching ^59^, are evaluated to guide users in their choice. In this preprint, we present an initial version of our evaluation of these methods, focusing on the quality control and the accuracy of their model using the default parameters. When comparing the maximum overall score over time, it is encouraging to see multiple incremental improvements over the state-of-the-art (**Supplementary Figure 15**). We believe that the benchmarking presented in this study will pave the way to the next series improvements in the field of trajectory inference.

In this study, we presented our first version of the evaluation of TI methods. We are convinced that the results we provided will be useful for both method users and tool developers, as we provide clear practical guidelines for users depending on their current knowledge of the trajectory, as well as an objective benchmark on which new methods can be tested. Nonetheless, our evaluation can be expanded on several points, all of which we will try to tackle in the near future:

- Inclusion of methods published during of after june 2017
- A parameter tuning for each method, both across all trajectory types, as well as on specific trajectory types.
- A test of robustness on noise, parameter changes and dataset size
- An evaluation of the stability of a method’s results when running the same parameter setting on the same dataset multiple times
- Evaluate the methods when given optional priors, and compare with performance without priors
- Assess the effect of noisy or incorrect prior information
- Test methods on datasets with no clear trajectory present
- Include feedback on the quality control scoring scheme, and update the scoring when method get updated
- Test the scalability of each method, both in the number of cells and the number of features
- Include more real datasets with complex trajectory types, as they become available
- Evaluate methods on other single-cell -omics datasets, such as proteomics and epigenomics data
- Provide functionality for visually interpreting and comparing predicted trajectories

Our evaluation indicated a large heterogeneity in the performance of the current TI methods, with Slingshot, TSCAN, and Monocle DDRTree, towering above all other methods. Nonetheless, we found that methods which did not perform well across all settings could still be useful in certain specific use cases. Indeed, on data containing more than one bifurcations, Monocle DDRTree clearly performed better than other methods. We found that this particular result was mainly caused by the fact that the default parameters of Monocle DDRTree preferably led to the detection of tree topologies, while those of Slingshot preferably found linear and bifurcation trajectories.

We managed to wrap the output of all methods into one common format. This not only allowed us to compare different methods with a gold standard, but could also be useful for TI users, as it allows the user to test multiple methods on the same data and compare the results without manual conversion of input and output. Furthermore, it makes it possible to directly compare the output of different methods, which opens up possibilities for new comparative visualisation techniques or ensemble methods. However, we acknowledge that our model has some limitations. Currently, It cannot take into account uncertainty of a cell’s position, which can both occur on the cellular ordering (e.g. when the position of a cell is uncertain within a branch) or on the trajectory topology (e.g. when the connections between branches are uncertain). Some current methods already model this uncertainty in some way, mainly on the cellular ordering^30^, and in the future we will adapt our output model to also allow this uncertainty.

The use of synthetic data to evaluate TI methods offers the advantage of having an exact gold standard to which the methods’ results can be compared. However, the use of synthetic data can also be questionable, because the model which generates the data does not necessarily reflect the intrinsic characteristics of true biological systems. This could bias an evaluation towards methods for which the underlying model best fits the model used for generating the data. Therefore, it is essential that results of an evaluation on synthetic data are confirmed using real data. In our study, the overall performance of methods was very similar between real and synthetic datasets, confirming the biological relevance of the synthetic data. Given this, we believe that our synthetic data can now be used to effectively prototype new TI methods for more complex trajectories such as disconnected graphs, for which the availability of real datasets is poor. Furthermore, it is expected that in the future, methods will be able to model even more complex cellular behaviors, such as multiple dynamic processes happening at parallel in a single cell, the integration of datasets from different patients or trajectories in a spatial context^1,4^. Synthetic data generated with our workflow could therefore be used to spearhead the development of these methods, given that currently only a limited number of real datasets are available for which the methods could be useful. Furthermore, we believe that our data generation workflow could also be used to evaluate other types of single-cell modelling techniques, such as single-cell network inference, clustering and normalisation. We sincerely hope that such efforts will lead to a more rapid development of accurate methods, and will in the near future provide a package which can be used to simulate synthetic data for a wide variety of single-cell modelling problems.

## Methods

### Trajectory inference methods

#### Method selection

We gathered a list of 59 trajectory inference methods (**Table 1**), by searching in literature for “trajectory inference” and “pseudotemporal ordering”, and based on two existing lists found online^74,76^. A continuously updated list can also be found online). We welcome any contributions by creating an issue at github.com/dynverse/dynverse/issues.

Methods were excluded from the evaluation based on several criteria: (a) Not free, (b) Unavailable, (c) Superseded by another method, (d) Requires data types other than expression, (e) No programming interface, (f) Unresolved errors during wrapping,(g)Too slow (requires more than one hour on a 100×100 dataset), (h) Doesn’t return an ordering, (i) Requires additional user input during the algorithm (not prior information), (j) Published later than 2017-05-01 to be included in the current version of the evaluation, (k) This method is not published in preprint or a peer-reviewed journal. This resulted in the inclusion of 29 methods in the evaluation (**Table 1**).

#### Method input

As input, we provided for each method either the raw count data (after cell and gene filtering) or normalised expression values, based on the description in the methods documentation or from the study describing the method. Furthermore, when required, we also provided a maximum of 7 types of prior information. This prior information was extracted from the gold/silver standards as follows:

- **Start cells** The identity of one or more start cells. For both real and synthetic data, a cell was chosen which was the closest (in geodesic distance) from each milestone with only outgoing edges. For ties, one random cell was chosen. For cyclic datasets, a random cell was chosen.
- **End cells** The identity of one or more end cells. Similar as the start cells, but now for every state with only ingoing edges.
- **# end states** Number of terminal states. Number of milestones with only ingoing edges.
- **Grouping** For each cell a label to which state/cluster/branch it belongs. For real data, the states from the gold/silver standard. For synthetic data, each milestone was seen as one group, and cells were assigned to their closest milestone.
- **# branches** Number of branches/intermediate states. For real data, the number of states in the gold/silver standard. For synthetic data, the number of milestones.
- **Time course** For each cell a time point from which it was sampled. If available, directly extracted from the gold standard. For synthetic data: four timepoints were chosen, at which the cells were “sampled” to provide a time course information reflecting the one provided in real experiments.

#### Common trajectory model

Due to the absence of a common format for trajectory models, most methods return a unique set of output formats with few overlap. We therefore created wrappers around each method (available at github.com/dynverse/dynmethods) and postprocessed its output into a common probabilistic trajectory model (**Supplementary Figure 16a**). This model consists of three parts.(i)The milestone network represents the overall network topology, and contains edges between different milestones and the length of the edge between them. (ii) The milestone percentages contain, for each cell, its position between milestones, and sums for each cell to one. (iii) The trajectory model also allows for regions of delayed commitment, where cells are allowed to be positioned between three or more connected milestones. Regions must be explicitly defined in the trajectory model. Per region, one milestone must be directly connected to all other milestones in the network.

Depending on the output of a method, we used different strategies to convert the output to our model (**Supplementary Figure 16b**). Special conversions are denoted by an *, and will be explained in more detail below.

- **Type 1, direct:** GPfates, reCAT, SCIMITAR, SLICE* and Wishbone. These methods assign each cell to a branch together with a pseudotime value and a branch network. The branch network is used as the milestone network, the percentages of a cell are proportional with its branch pseudotime.
- **Type 2, linear pseudotime:** Embeddr, Ouija, Ouijaflow, Phenopath, Pseudogp, SCORPIUS, Topslam, Wanderlust and Waterfall. These methods return a pseudotime value for each cell. The milestone network will consist of a single edge between two milestones, where cells are positioned on the transition proportional to their pseudotime value.
- **Type 3, end state probability:** MFA* and SCOUP. These methods return a global pseudotime value and a probability for every end state. We use a single start state and add an edge to every end milestone each representing an end state. The global pseudotime then determines the distance from the begin milestone, the rest of the cell’s position is calculated by distributing the residual percentage over the end states, proportionally to the end state probabilities.
- **Type 4, cluster assignment:** Mpath and SCUBA. These methods return a cluster assignment and a cluster network. The cluster network was used as milestone network, each cell received percentage 1 or 0 based on its cluster assignment.
- **Type 5, projection onto nearest branch:** DPT*, Slingshot, StemID and TSCAN. These methods returning a cluster assignment, cluster network and dimensionality reduction. We projected each cell on the closest point on the edges between its own cluster and neighbouring clusters within the dimensionality reduction. This projection allowed us to give each cell a pseudotime within the edge, which was then converted into our model as described above, using the cluster network as milestone network.
- **Type 6, cell graph:** cellTree Gibbs, cellTree Maptpx, cellTree VEM, Monocle DDRTree, Monocle ICA, Sincell* and SLICER. These methods return a network of connected cells, and determine which cells are part of the “backbone”. One milestone is created for each cell that is part of the backbone and has a degree ≠ 2. Cells are positioned on the closest segment that is part of the backbone.

Special conversions were necessary for certain methods:

- **DPT** We projected the cells onto the cluster network, consisting of a central milestone (this cluster contains the cells which were assigned to the “unknown” branch) and three terminal milestones, each corresponding to a tip point. This was then processed as described above.
- **Sincell** To constrain the number of milestones this method creates, we merged two cell clusters iteratively until the percentage of leaf nodes was below a certain cutoff, default at 25%. This was then processed as described above.
- **SLICE** As discussed in the vignette of SLICE, we ran principal curves one by one for every edge detected by SLICE. This local pseudotime was then processed as above.
- **MFA** We used the branch assignment as state probabilities, which together with the global pseudotime were processed as described above.

### Real datasets

We gathered a list of real datasets by searching for “single-cell” at the Gene Expression Omnibus and selecting those datasets in which the cells are sampled from different stages in a dynamic process (**Supplementary Table 1** and **Supplementary Figure 4**). The scripts to download and process these datasets will be made available on our repository (github.com/dynverse/dynalysis). Whenever possible, we preferred to start from the raw counts data. These raw counts were all normalised and filtered using a common pipeline, discussed later. We determined a reference standard for every dataset using labelling provided by the author’s, and classified the standards into gold and silver based on whether this labelling was determined by the expert using the expression (silver standard) or using other external information (such as FACS or the origin of the sample, gold standard) (**Supplementary Table 1**).

### Synthetic datasets

Our workflow to generate synthetic data is based on the well established workflow used in the evaluation of network inference methods^70,77^ and consists of four main steps: network generation, simulation, gold standard extraction and simulation of the scRNA-seq experiment (**Supplementary Figure 5**). At every step, we took great care to mimic real cellular regulatory networks as best as possible, while keeping the model simple and easily extendable. For every synthetic dataset, we used a random real dataset as a reference dataset (from those described earlier), making sure the number of variable genes and cells were similar.

### Network generation

One of the main processes involved in cellular dynamic processes is gene regulation, where regulatory cascades and feedback loops lead to progressive changes in expression and decision making. The exact way a cell choses a certain path during its differentiation is still an active research field, although certain models have already emerged and been tested in vivo. One driver of bifurcation seems to be mutual antagonism, where genes^78^ strongly repress each other, forcing one of the two to become inactive^79^. Such mutual antagonism can be modelled and simulated^80,81^. Although such a two-gene model is simple and elegant, the reality is frequently more complex, with multiple genes (grouped into models) repressing each other^82^.

To simulate certain trajectory topologies, we therefore designed module networks in which the cells follow a particular trajectory topology given certain parameters (**Supplementary Figure 17**). Two module networks generated linear trajectories (linear and linear long), two generated simple forks (bifurcating and converging), one generated a complex fork (trifurcating), one generated a rooted tree (consecutive bifurcating) and two generated simple undirected graphs (bifurcating loop and bifurcating convergence).

From these module networks we generated gene regulatory networks in two steps: the main regulatory network was first generated, and extra target genes from real regulatory networks were added. For each dataset, we used the same number of genes as were differentially expressed in the real datasets. 5% of the genes were assigned to be part of the main regulatory network, and were randomly distributed among all modules (with at least one gene per module). We sampled edges between these individual genes (according to the module network) using a uniform distribution between 1 and the number of possible targets in each module. To add additional target genes to the network, we assigned every regulator from the network to a real regulator in a real network (from regulatory circuits^83^), and extracted for every regulator a local network around it using personalized pagerank (with damping factor set to 0.1), as implemented in the `page_rank` function of the *igraph* package.

### Simulation of gene regulatory systems using thermodynamic models

To simulate the gene regulatory network, we used a system of differential equations similar to those used in evaluations of gene regulatory network inference methods^77^. In this model, the changes in gene expression (*x*_*i*_) and protein expression (*y*_*i*_) are modeled using ordinary differential equations^70^ (ODEs):

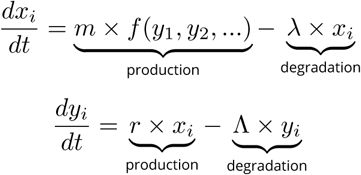

where *m, λ, r* and Λ represent production and degradation rates, the ratio of which determines the maximal gene and protein expression. The two types of equations are coupled because the production of protein *y*_*i*_ depends on the amount of gene expression *x*_*i*_, which in turn depends on the amount of other proteins through the activation function *f* (*y*_1_, *y*_2_, *…*).

The activation function is inspired by a thermodynamic model of gene regulation, in which the promoter of a gene can be bound or unbound by a set of transcription factors, each representing a certain state of the promoter. Each state is linked with a relative activation *α*_*j*_, a number between 0 and 1 representing the activity of the promoter at this particular state. The production rate of the gene is calculated by combining the probabilities of the promoter being in each state with the relative activation:

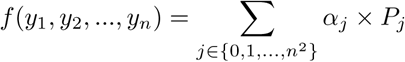

The probability of being in a state is based on the thermodynamics of transcription factor binding. When only one transcription factor is bound in a state:

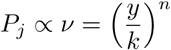

Where the hill coefficient *n* represents the cooperativity of binding and *k* the transcription factor concentration at half-maximal binding. When multiple regulators are bound:

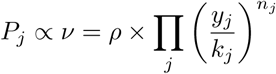

where *ρ* represents the cooperativity of binding between the different transcription factors.

*P*_*i*_ is only proportional to *v* because *v* is normalized such that ∑ _*i*_ *P*_*i*_ = 1

To each differential equation, we added an additional stochastic term:

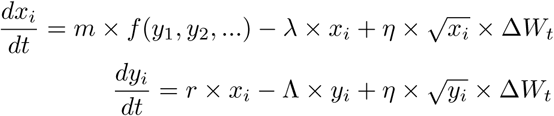

with Δ*W*_*t*_ ∼ 𝒩 (0, *h*).

Similar to^70^, we sample the different parameters from random distributions, given in **Supplementary Table 4**.

We converted each ODE to an SDE by adding a chemical Langevin equation, as described in^70^. These SDEs were simulated using the Euler–Maruyama approximation, with time-step *h* = 0.01 and noise strength *η* = 8. The total simulation time varied between 5 for linear and bifurcating datasets, 10 for consecutive bifurcating, trifurcating and converging datasets, 15 for bifurcating converging datasets and 30 for linear long, cycle and bifurcating loop datasets. The burn-in period was for each simulation 2. Each network was simulated 32 times.

### Simulation of the single-cell RNA-seq experiment

For each dataset we sampled the same number of cells as were present in the reference real dataset, limited to the simulation steps after burn-in. Next, we used the Splatter package^84^ to estimate the different characteristics of a real dataset, such as the distributions of average gene expression, library sizes and dropout probabilities. We used Splatter to simulate the expression levels *λ*_*i,j*_ of housekeeping genes *i* (to match the number of genes in the reference dataset) in every cell *j*. These were combined with the expression levels of the genes simulated within a trajectory. Next, true counts were simulated using 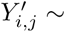 Poi(*λ*_*i,j*_). Finally, we simulated dropouts by setting true counts to zero by sampling from a Bernoulli distribution using a dropout probability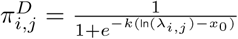.

### Gold standard extraction

Because each cellular simulation follows the trajectory at its own speed, knowing the exact position of a cell within the trajectory topology is not straightforward. Furthermore, the speed at which simulated cells make a decision between two or more alternative paths is highly variable. To estimate a cell’s position during a simulation within the trajectory topology, we therefore used the known progression of the modules, given in **Supplementary Figure 17c**, as a backbone. We smoothed the expression in each simulation using a rolling mean with a window of 50 time steps, and then calculated the average module expression along the simulation. We used dynamic time warping, implemented in the dtw R package^85,86^, with an open end to align a simulation to all possible module progressions, and then picked the alignment which minimised the normalised distance between the simulation and the backbone. In case of cyclical trajectory topologies, the number of possible milestones a backbone could progress through was limited to 20.

### Expression normalisation pipeline

We used a standard single-cell RNA-seq preprocessing pipeline which applies parts of the scran and scater Bioconductor packages^87^. The advantages of this pipeline is that it works both with and without spike-ins, and includes a harsh cell filtering which looks at abnormalities in library sizes, mitochondrial gene expression, and number of genes expressed using median absolute deviations (set to 3). We required that a gene was expressed in at least 5% of the cells, and that it should have an average expression higher than 0.05. Furthermore, we used the pipeline to select the most highly variable genes, using a false discovery rate of 5% and a biological component higher than 0.5. As a final filter, we removed both zero genes and cells until convergence.

### Evaluation metrics

The importance of using multiple metrics to compare complex models has been stated repeatedly^88^. We defined three metrics for comparing the likeness of predicted trajectories to a gold standard (**Figure 1c**). Each metric assesses the performance of a different aspect of the trajectory (**Supplementary Figure 18**); (i) the correlation metric measures the similarity in pairwise cell-cell distances; (ii) the RF MSE metric assesses whether cell neighbourhoods are similar in both trajectories; and (iii) the edge flip scores assesses similarity in trajectory topologies.

### Correlation between geodesic distances

The similarity in cell ordering between two trajectories is assessed by calculating the geodesic distances between each pair of cells for both trajectories. The definition of a geodesic distance between two cells part of the common trajectory model will be demonstrated using a toy example (**Supplementary Figure 19**).

The geodesic distance between two cells depends on whether they are (i) on the same transition, or (ii) in different transitions. In the first case, the distance is defined as the product of the difference in milestone percentages and the length of the transition they both reside on. For cells *a* and *b* in the example, *d*(*a, b*) is equal to 1 × (0.9 − 0.2) = 0.7. In the latter case, the distances between the cells and all of their neighbouring milestones will be calculated. These distances in combination with the milestone network are used to calculate the shortest path distance between the two cells. For cells *a* and *c* in the example, *d*(*a, X*) = 1 *×* 0.9 and *d*(*c, X*) = 3 *×* 0.2, and therefore *d*(*a, c*) = 1 *×* 0.9 + 3 *×* 0.2.

According to the defined common trajectory model (**Figure 1b**), cells are also allowed to have a delayed commitment. In a region of delayed commitment, one milestone will be connected to all other milestones as per the milestone network. In this case, the distance between two cells both inside a region of delayed commitment is calculated as the manhattan distances between the milestone percentages weighted by the transition weights from the milestone network. For cells *d* and *e* in the example, *d*(*d, e*) is equal to 0 (0.3 0.2) + 2 (0.7 0.2) + 3 (0.4 0.1), which is equal to 1.9. The distance between two cells where one is part of a region of delayed commitment is calculated similarly to the previous paragraph, by first calculating the between the cells and their neighbouring milestones first, then calculating the shortest path distances between the two.

Finally, calculating all pairwise distances between cells would scale poorly for trajectories with large numbers of cells. For this reason, a set of waypoint cells are defined *a priori*, and only the distances between the waypoint cells and all other cells is calculated, in order to calculate the correlation of geodesic distances of two trajectories. The waypoints are determined by viewing each milestone, transition and region of delayed commitment as a collection of cells, and sampling cells from the different collections weighted by the total number of cells within that collection. For calculating the correlation of geodesic distances between two trajectories, the distances between all cells and the union of both waypoint sets is computed. For the benchmark evaluation, the total number of waypoints sampled from a trajectory was one hundred.

### Random forest prediction error

Although the correlation between geodesic distances directly assesses the position of the cells in the trajectory, a bad correlation does not directly imply that similar cells were not grouped together by the method, as illustrated in **Supplementary Figure 18b,c**. For example, certain methods will inevitably reach an incorrect ordering because they cannot handle the correct trajectory type, but these methods could still correctly place similar cells next to each other. We therefore also included a metric which looks at the local neighbourhood of each cell, and assesses whether this neighbourhood can accurately predict the position of this cell in the gold standard. We used a Random Forest regression, implemented in the *ranger* package^89^ to separately predict milestone percentages of every cell in the gold standard, using the milestone percentages of these cells in the prediction as features. We then used the out-of-bag mean-squared error on these percentages to score each method’s capability of predicting the correct neighbourhood of each cell.

### Edge flip score

As a third independent score, we assessed the similarity between the milestone network topologies. We first simplified each network, by merging consecutive linear edges into one edge, and adding new milestones within self loops such that *A → A* would be converted *A → B → C → D*, by adding an intermediate node to linear networks. Because we are interested in the overall similarity between two topologies irrespective of the direction of the edges, the network was made undirected. Next, we define the edge flip score as the minimal number of edges which should be added or removed to convert one network into the other, divided by the total number of edges in both networks. This problem is equivalent to the maximum common edge subgraph problem, a known NP-hard problem without a scalable solution^90^. We implemented a branch and bound approach for this problem, by first enumerating all possible edge additions and removals with the minimal number of edges (the edge difference between the two networks) and if none of the new networks was isomorphic, we tried out all solutions with additional two edge changes. To further limit the search space, we made sure the degree distributions between the two networks were similar, before assessing whether the two networks were isomorphic using the BLISS algorithm^91^, as implemented in the R *igraph* package.

A comparison of edge flip scores between common trajectory topologies is illustrated in **Supplementary Figure 20**.

### Aggregation of scores

**Supplementary Figure 21** illustrates the full set of aggregations performed on the raw scores in order to arrive at the final ranking of methods. In order to make the scores produced by the different metrics comparable to one another, across datasets of varying difficulty, the raw scores are percentile rank transformed per metric per dataset to a [0, 1] range. The arithmetic mean is calculated across replicates. In order to ensure a method only obtains a high score if it scores well on all three metrics, the harmonic mean across metrics is calculated. At this point, the aggregated scores of all methods across all datasets is combined. As there can be an overrepresentation of datasets of a certain trajectory type, first an arithmetic mean is calculated per trajectory type, followed by an overall arithmetic mean across all trajectory types, thus obtaining a ranking of the methods.

### Benchmark

#### Method execution

Each execution of a method on a dataset was performed in a separate task as part of a gridengine job. Each task was allocated one CPU core of an Intel(R) Xeon(R) CPU E5-2665 0 @ 2.40GHz, and was provided a maximum of 32GB in memory and 6 hours of wall time. One R session was started for each task, with the environment variable R_MAX_NUM_DLLS set to 500. The base::set.seed was overridden in order to prevent stochastic TI methods from pretending to be deterministic and/or robust. Timings of methods were measured for different steps along the executions, including preprocessing, postprocessing, each of the different metrics, and the method itself.

#### Effect of prior information

To assess the effect of prior information on the performance of a method, we compared the performance of methods which do or do not require the prior information using a two-tailed Mann–Whitney U test. P-values were controlled for multiple testing using Benjamini-Hochberg correction.

#### Effect of algorithm components

To assess the effect of algorithm components on the performance of a method, we used (1) a two-tailed Mann–Whitney U test, as implemented in the wilcox.test R function, and (2) increase in node purity importance scores using random forest classification, as implemented in the R randomForest package, predicting whether a method scored better than the median score. To evaluate whether a method increased or decreased performance, we used the estimate of the location parameter of the Mann–Whitney U test. We only investigated algorithm components which were part of at least 4 different methods in our evaluation study. P-values were controlled for multiple testing using Benjamini-Hochberg correction.

### Method quality control

We created a transparent scoring scheme to check the quality of each method (**Supplementary Table 3**), based on several existing tool quality and programming guidelines in literature and online ^92,71,93,94,72,95,96,97,98,99,100,101,102,103,104,105^. The goal of this quality control in the first place is to stimulate the improvement of current methods, and the development of user and developer friendly new methods. The quality control assessed 6 categories, each looking at several aspects, which are further divided into individual items. The availability category checks whether the method is easily available, whether the code and dependencies can be easily installed, and how the method can be used. The code quality assesses the quality of the code both from a user perspective (function naming, dummy proofing and availability of plotting functions) and a developer perspective (consistent style and code duplication). The code assurance category is frequently overlooked, and checks for code testing, continuous integration^99^ and an active support system. The documentation category checks the quality of the documentation, both externally (tutorials and function documentation) and internally (inline documentation). The behaviour category assesses the ease by which the method can be run, by looking for unexpected output files and messages, prior information and how easy the trajectory model can be extracted from the output. Finally, we also assessed certain aspects of the study in which the method was proposed, such as publication in a peer-reviewed journal, the number of dataset in which the usefulness of the method was shown, and the scope of method evaluation in the paper.

Each aspect was further assigned to one or more applications, based on whether it influenced the user friendliness of the tool (such as code availability, good documentation or contains plotting functions), the developer friendliness of the tool (such as unit testing, inline documentation and a clear licensing), or indications that the tool will be broadly applicable on new datasets (such as being open-source, containing a good tutorial and support system, and being thoroughly evaluated in the study where it was published).

Each quality aspect received a weight depending on how frequently it was found in several papers and online sources which discuss tool quality (**Table 1**). This was to make sure that more important aspects, such as the open source availability of the method, outweighed other aspects, such as the availability of a graphical user interface. Within each aspect, we also assigned a weight to the individual questions being investigated (**Table 1**). For calculating the final score, we weighed each of the six categories equally.

### Trajectory types

We classified all possible trajectory topologies into distinct trajectory types, based on topological criteria. These trajectory types start from the most general trajectory type, a disconnected directed graph, and move down (within a directed acyclic graph structure), progressively becoming more simple until the two basic types: linear and cyclical (**Supplementary Figure 22a**). For every directed trajectory type, a corresponding undirected trajectory type can also be defined (**Supplementary Figure 22b**). In most cases, a method which was able to detect a complex trajectory type, was also able to detect less complex trajectory types, with the only exception being DPT (limited to bifurcating trajectories).

## Supplementary Material

## Supplementary Figures

**Supplementary Figure 1.**
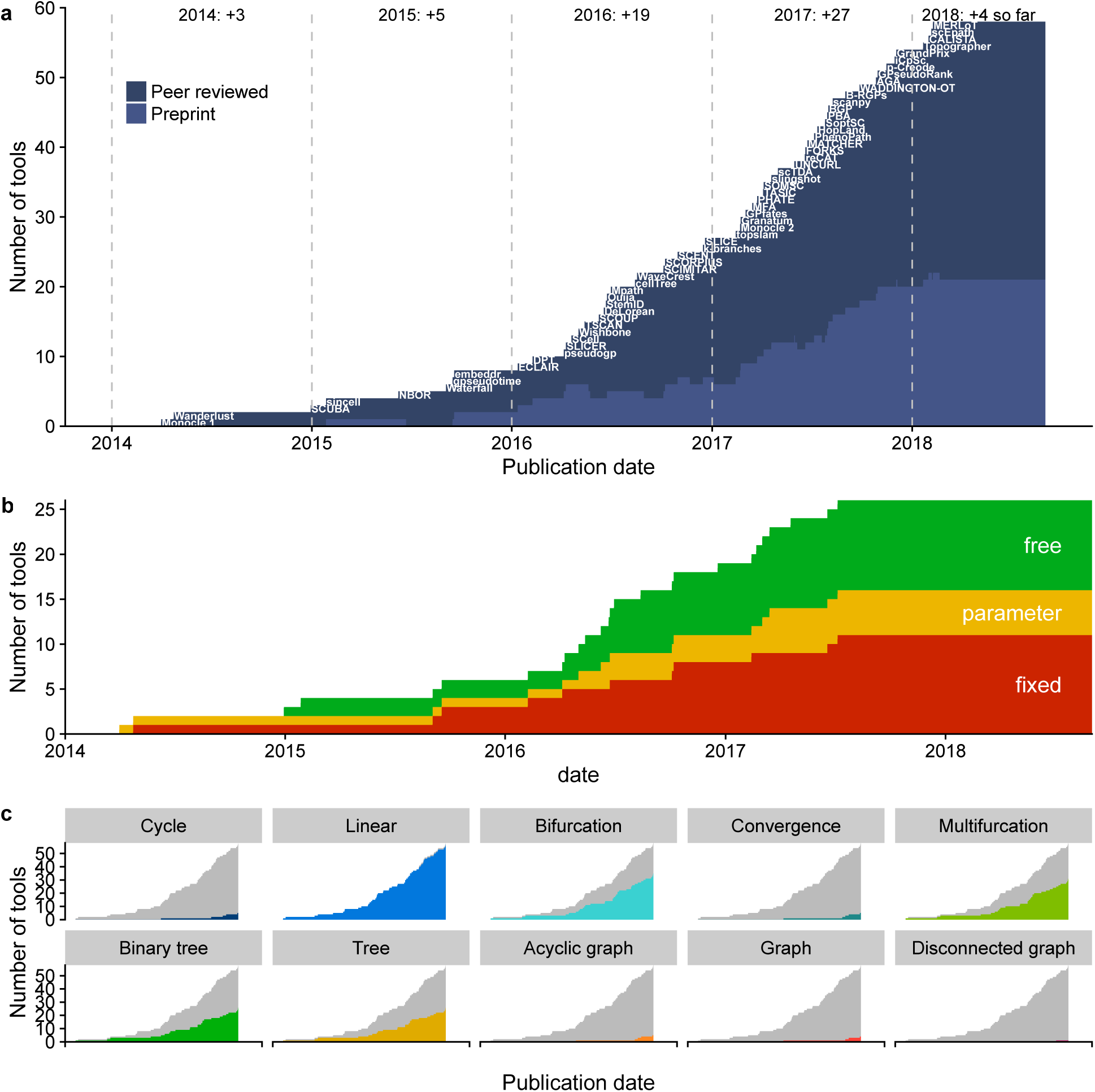
New TI methods over time. a) Number of methods published or in preprint. b) Number of methods fixing the trajectory topology, either by design (fixed) or through user parameters (parameter). c) Number of methods which can handle a particular type of trajectory.

**Supplementary Figure 2.**
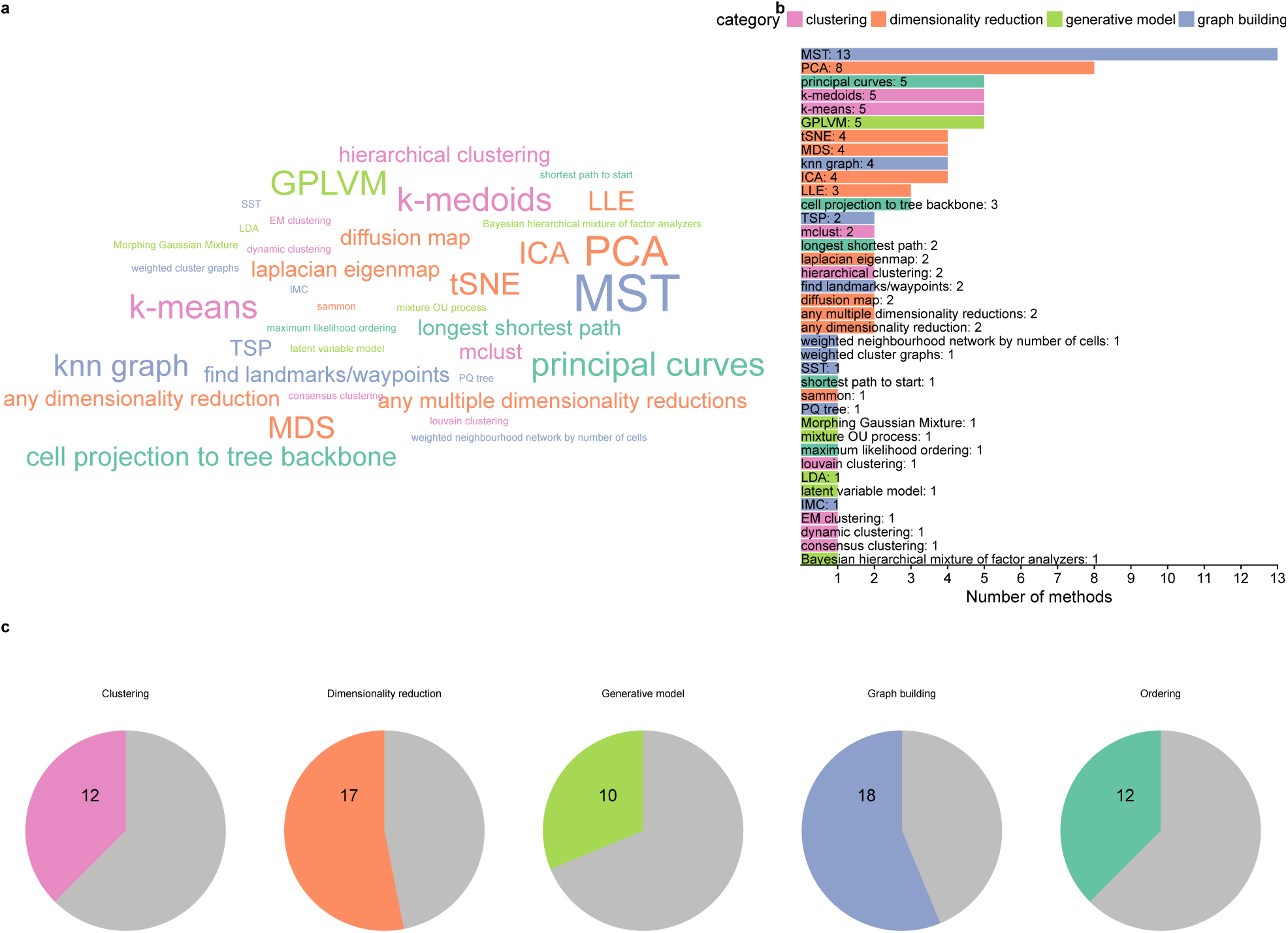
Common algorithmic components shared by different TI methods. Most components can be categorised into 4 categories: clustering (pink), dimensionality reduction (orange), generative models (green) and graph building (blue). a) Wordcloud of the common components. b) Number of times a component was shared between methods. c) Number of methods containing a particular category.

**Supplementary Figure 3.**
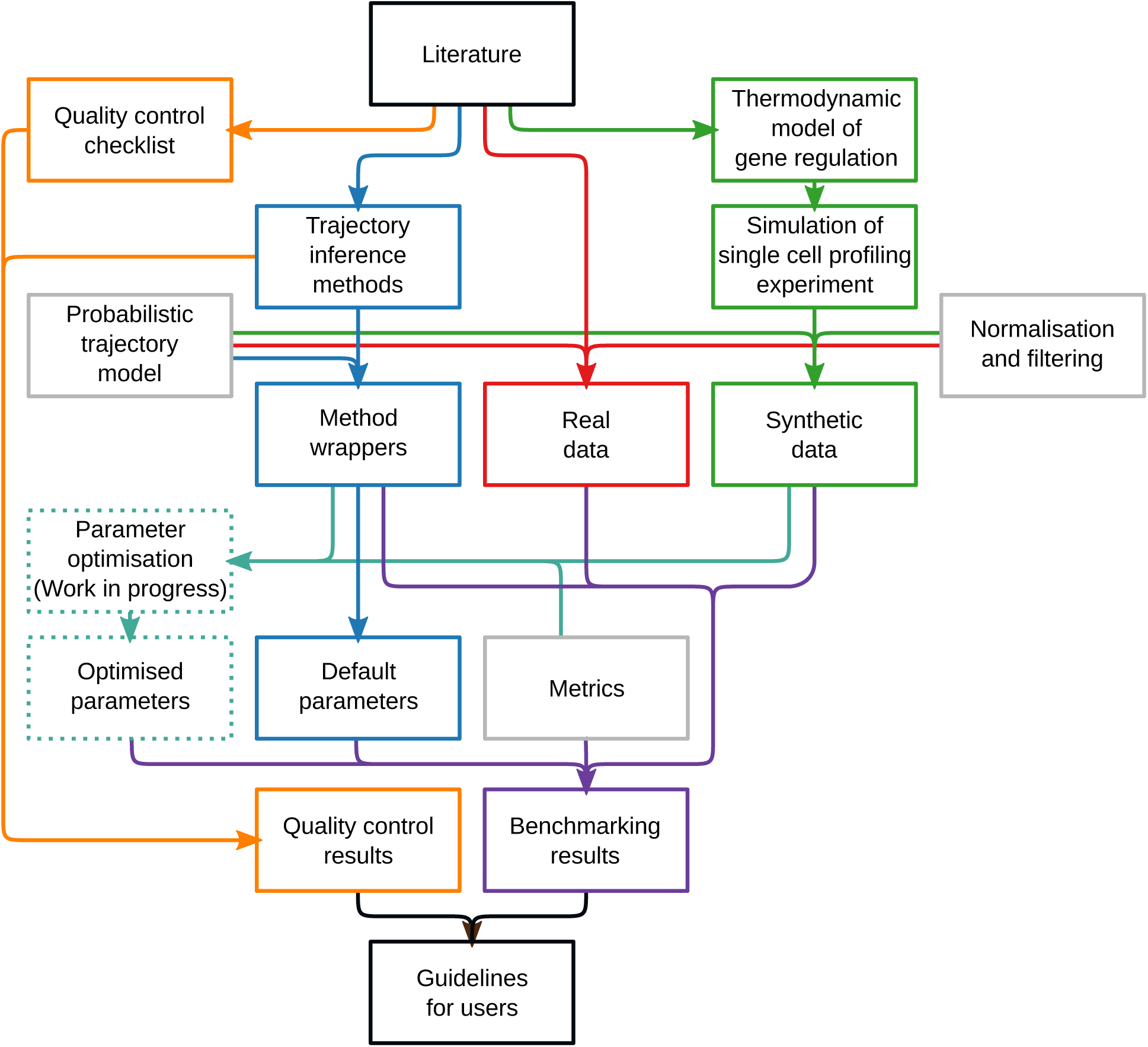
Extended overview of our evaluation pipeline. From literature we extracted a quality control checklist, a list of trajectory inference methods, a set of real datasets containing a trajectory and real regulatory networks used to generate the synthetic data. We created a wrapper of each method, so that its output was transformed into a common probabilistic trajectory model, and used these to infer trajectories on all real and synthetic datasets using their default parameters. Using several similarity metrics, we compared the gold standard of the real and synthetic datasets with the inferred trajectories. We also evaluated the quality of each method using the quality control checklist, and used both the benchmark results and quality control results to produce a final set of guidelines for method’s users.

**Supplementary Figure 4.**
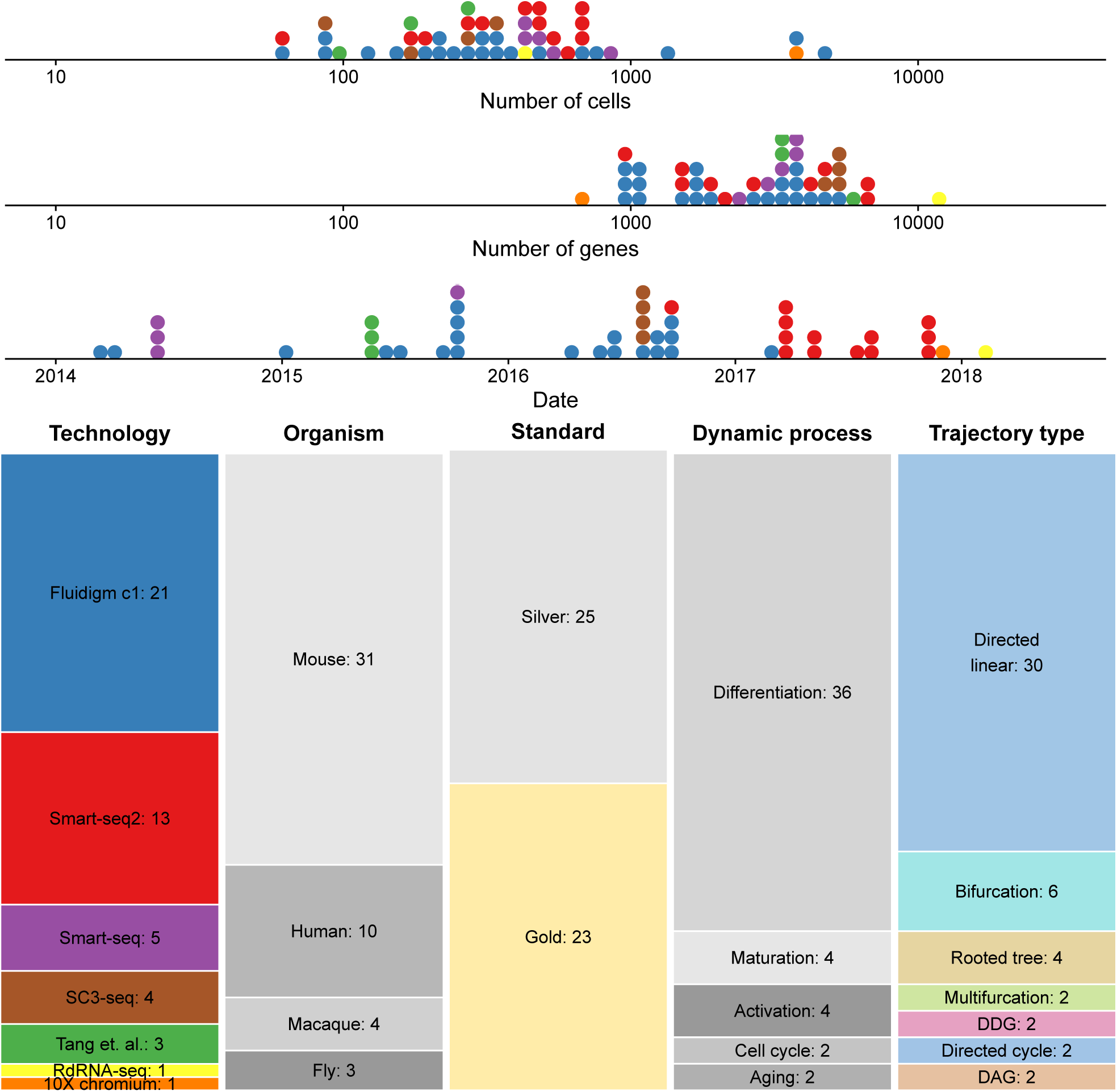
Characteristics of the real datasets used for the evaluation. Top: distributions of number of cells and genes (after filtering) and the date these datasets were published. Bottom: Diversity of technologies which were used to generate the dataset, the organism, whether we extracted a silver or a gold standard from the data, the type of dynamic process and the trajectory type present in the data.

**Supplementary Figure 5.**
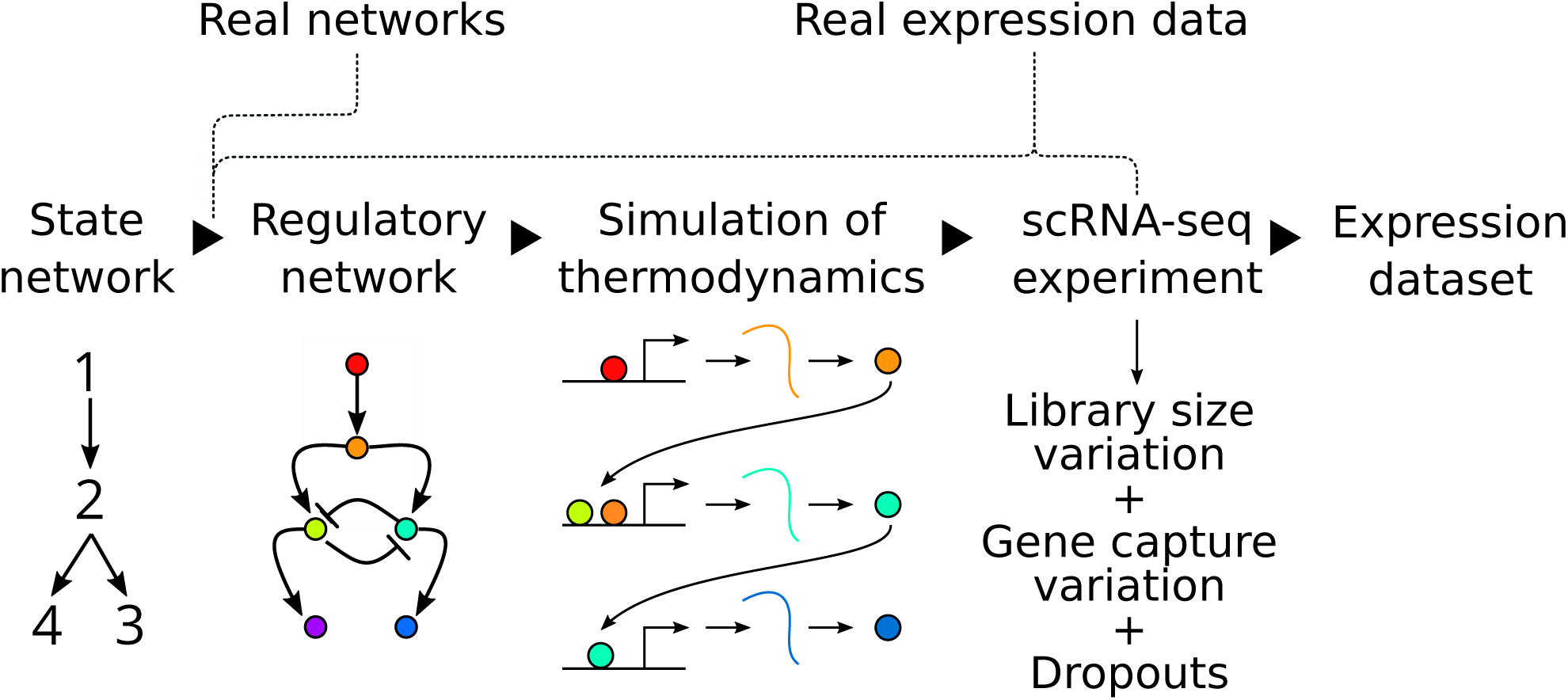
Workflow to generate the synthetic data. Starting from a state network, we extracted a regulatory network using real networks which is expected to induce such a set of state transitions when simulated. We simulated this network using a detailed model of gene regulation which includes the thermodynamics of transcription factor binding. Finally, we simulated the single-cell RNA-seq (scRNA-seq) experiment itself, by matching the distribution of expression and dropouts to a real reference dataset.

**Supplementary Figure 6.**
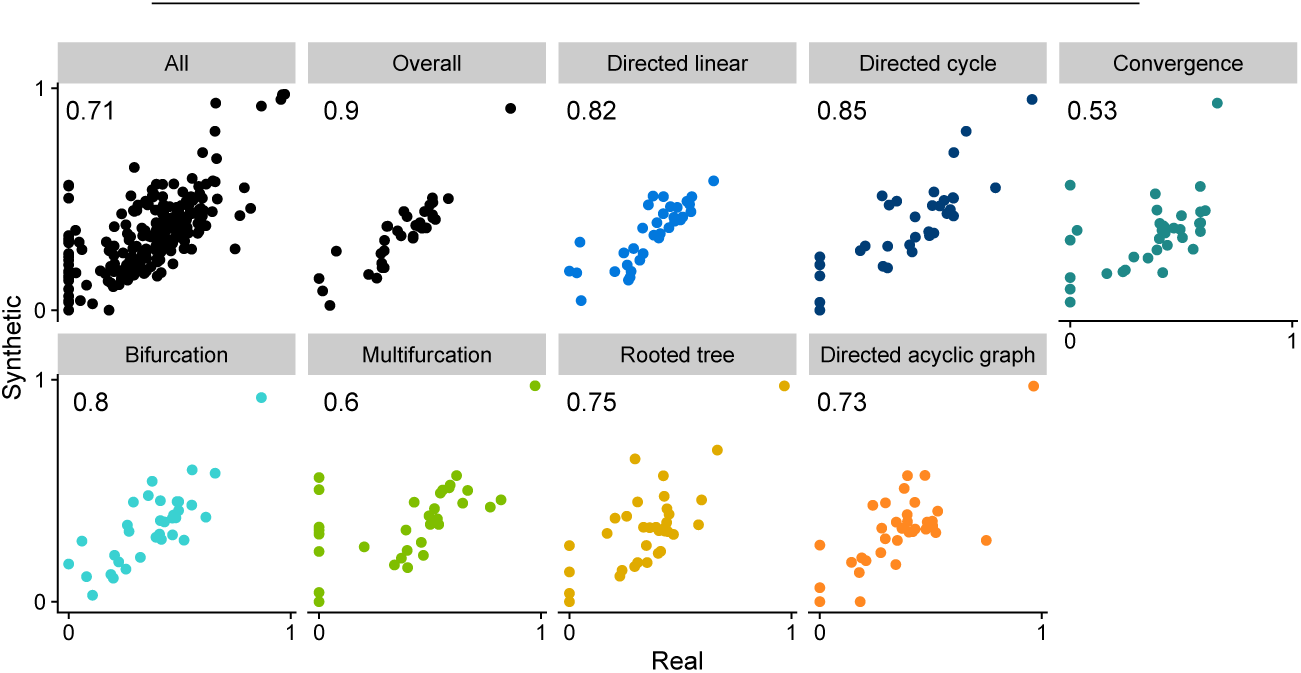
Comparison between performance on real and synthetic datasets across trajectory types. Given in the top-left corner of each panel is the Pearson’s correlation.

**Supplementary Figure 7.**
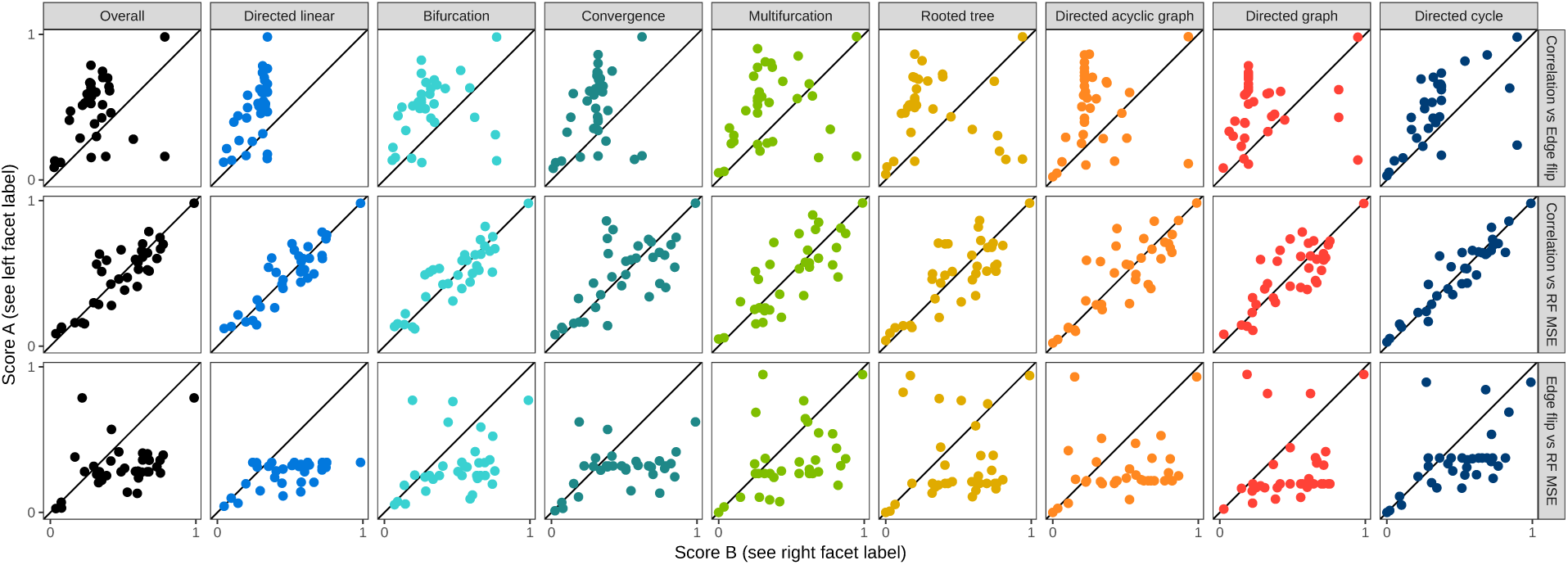
Comparison between the different metrics across trajectory types. For each row, a different pairwise comparison is given between two metrics.

**Supplementary Figure 8.**
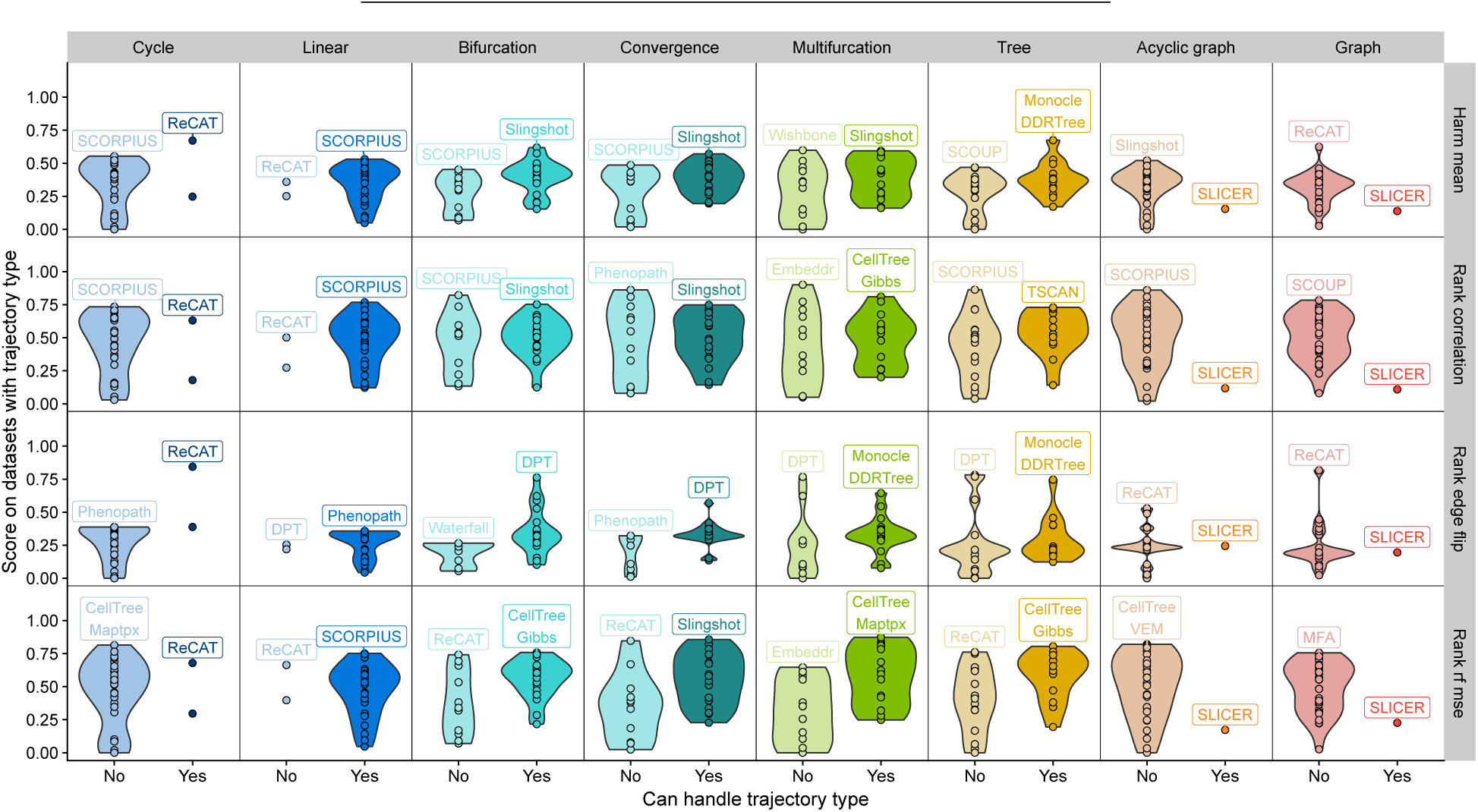
Comparison between the performance of methods able to handle a trajectory type with those who don’t. Shown is the performance on those datasets containing a particular trajectory type (top), according to several metrics (right).

**Supplementary Figure 9.**
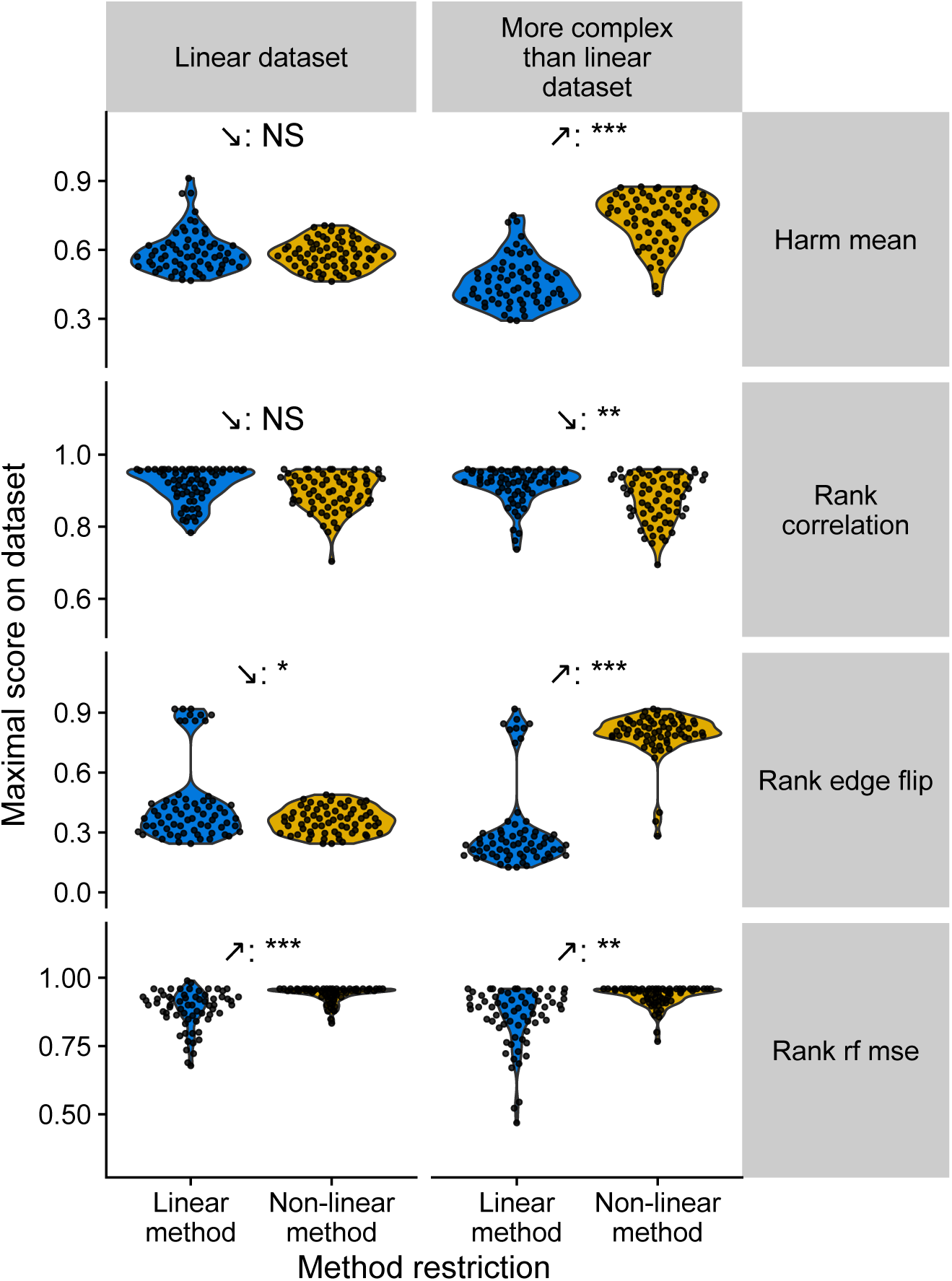
Comparison of the performance between linear and non-linear methods. Methods were split in linear methods (those which can either only detect cyclic or linear trajectories) and non-linear methods. Their performance is shown on linear and non-linear datasets according to several performance metrics (right). NS: Not significant, *: p-value < 0.05, **: p-value < 0.001, ***: p-value < 0.0001

**Supplementary Figure 10.**
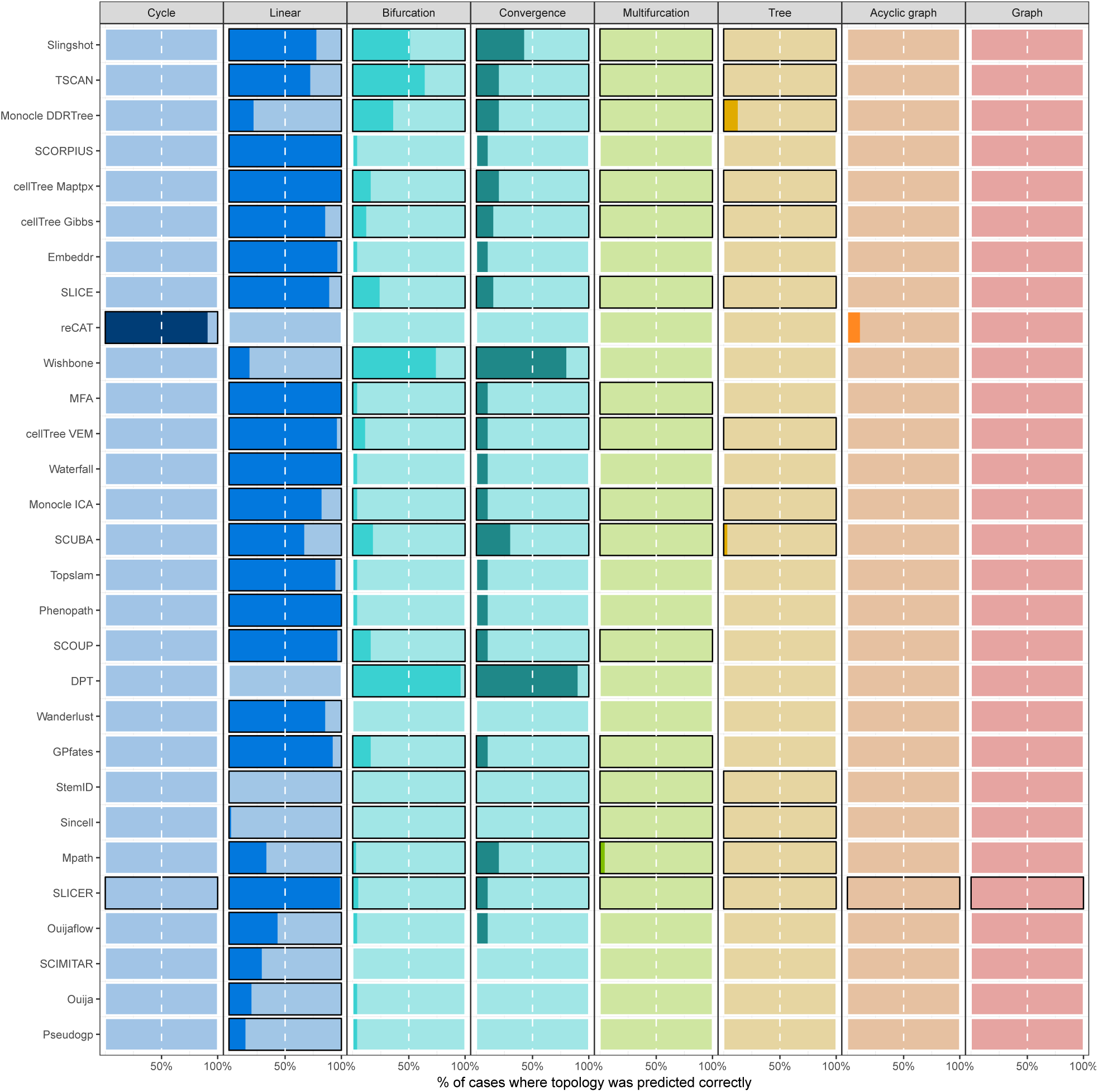
Sensitivity of each TI method in predicting the correct trajectory topology. Shown are the % of datasets where a method is able to predict the exact correct trajectory topology, depending on the trajectory type present in the data. When a method is able to predict a particular trajectory type, the box is surrounded by a black border.

**Supplementary Figure 11.**
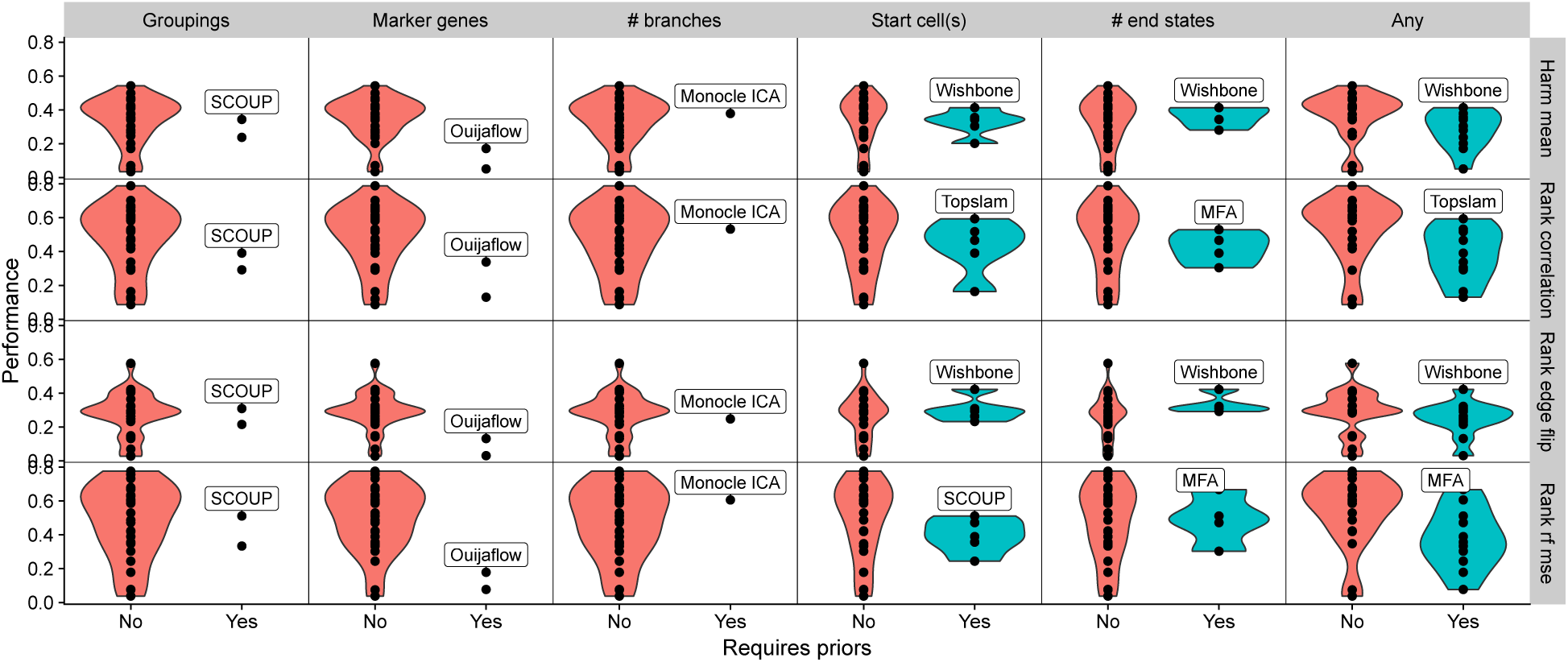
Comparison of method performance depending on whether the method required a certain prior or not. The top performing method which requires a particular type of prior information is labelled.

**Supplementary Figure 12.**
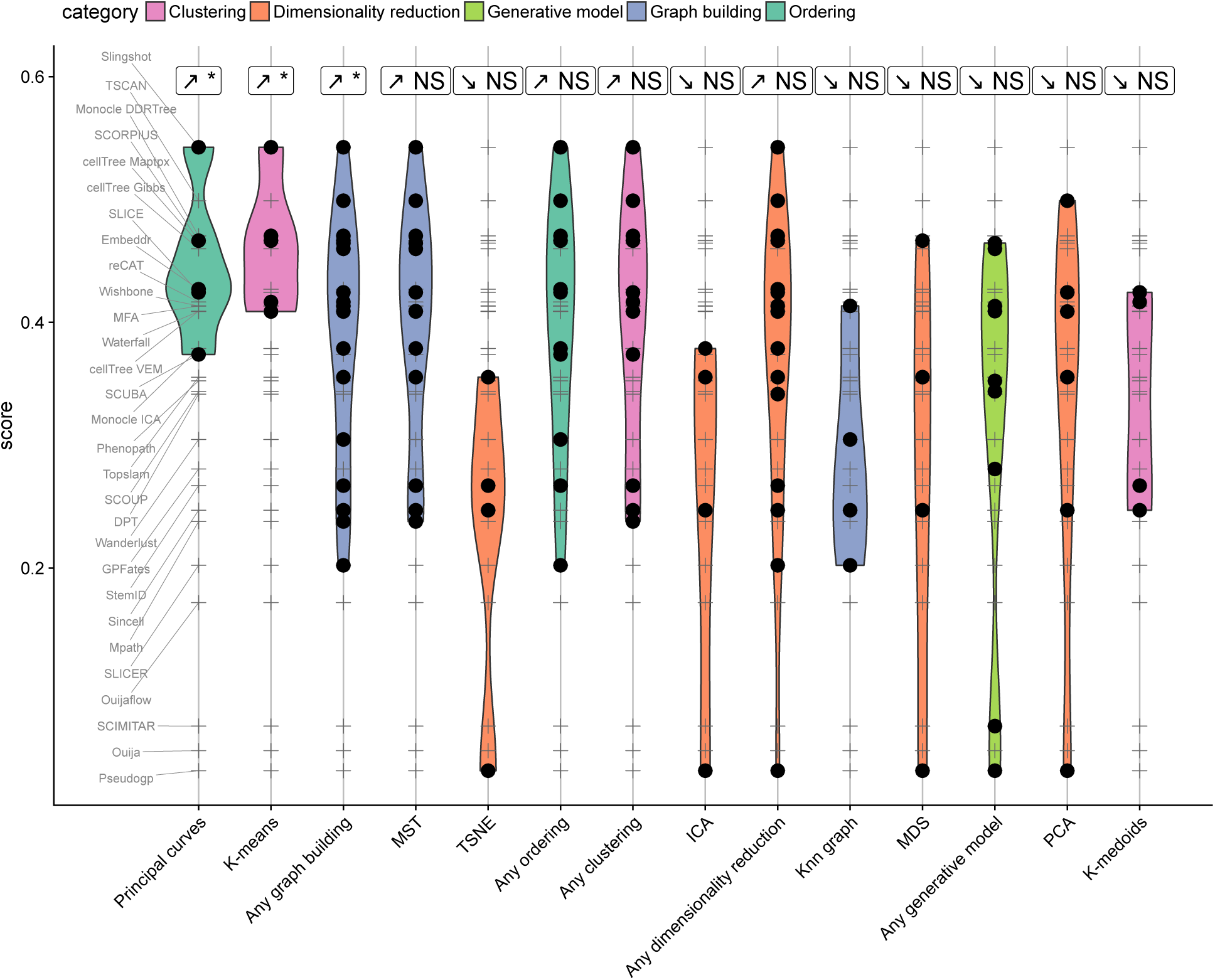
Score of methods containing particular algorithmic components. Shown are the algorithmic components present in four or more methods, and whether the performance of methods containing a certain component was significantly higher than all the other methods. NS: Not significant. *: corrected p-value < 0.05

**Supplementary Figure 13.**
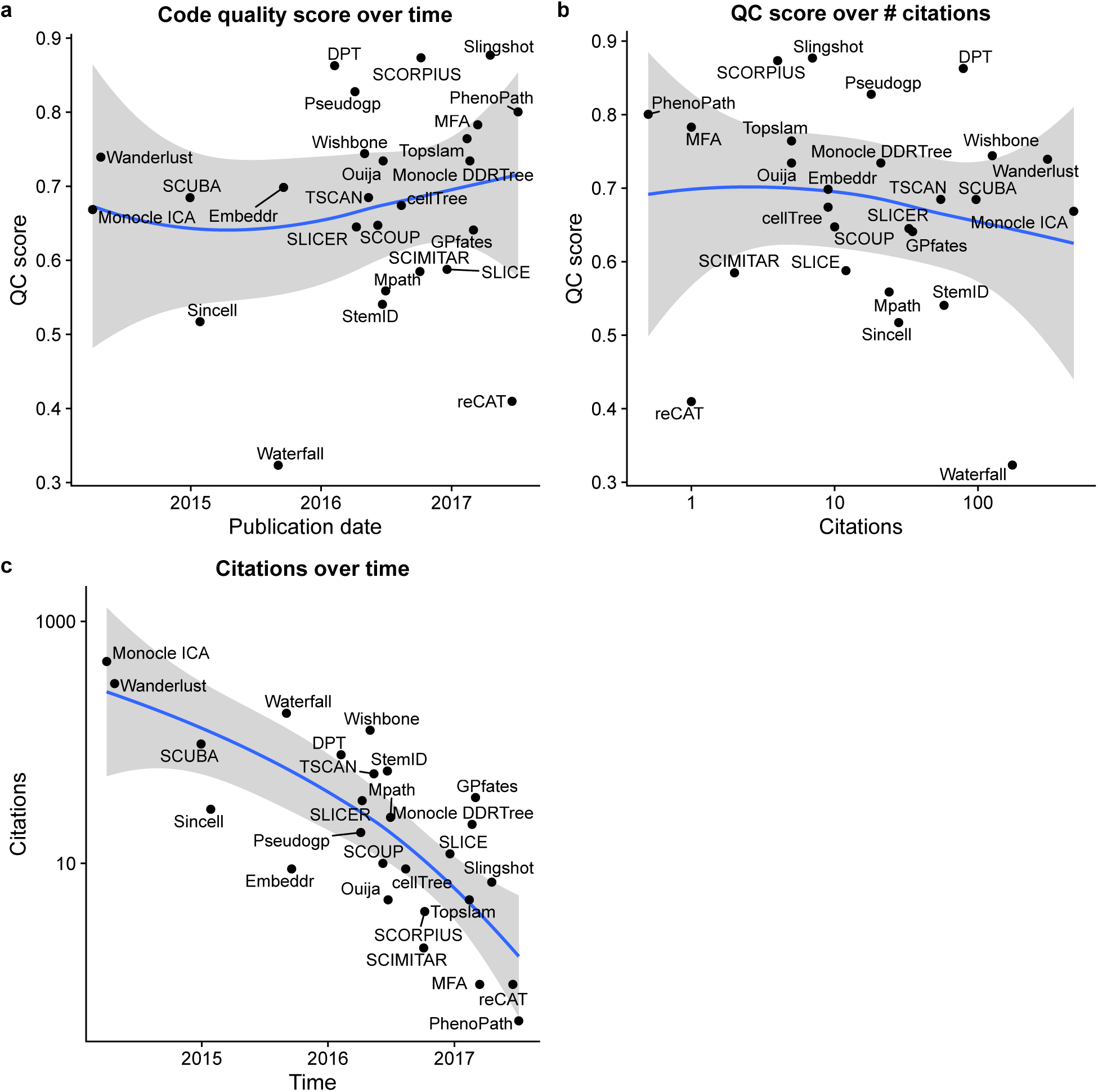
Quality control scores and number of citations over time.

**Supplementary Figure 14.**
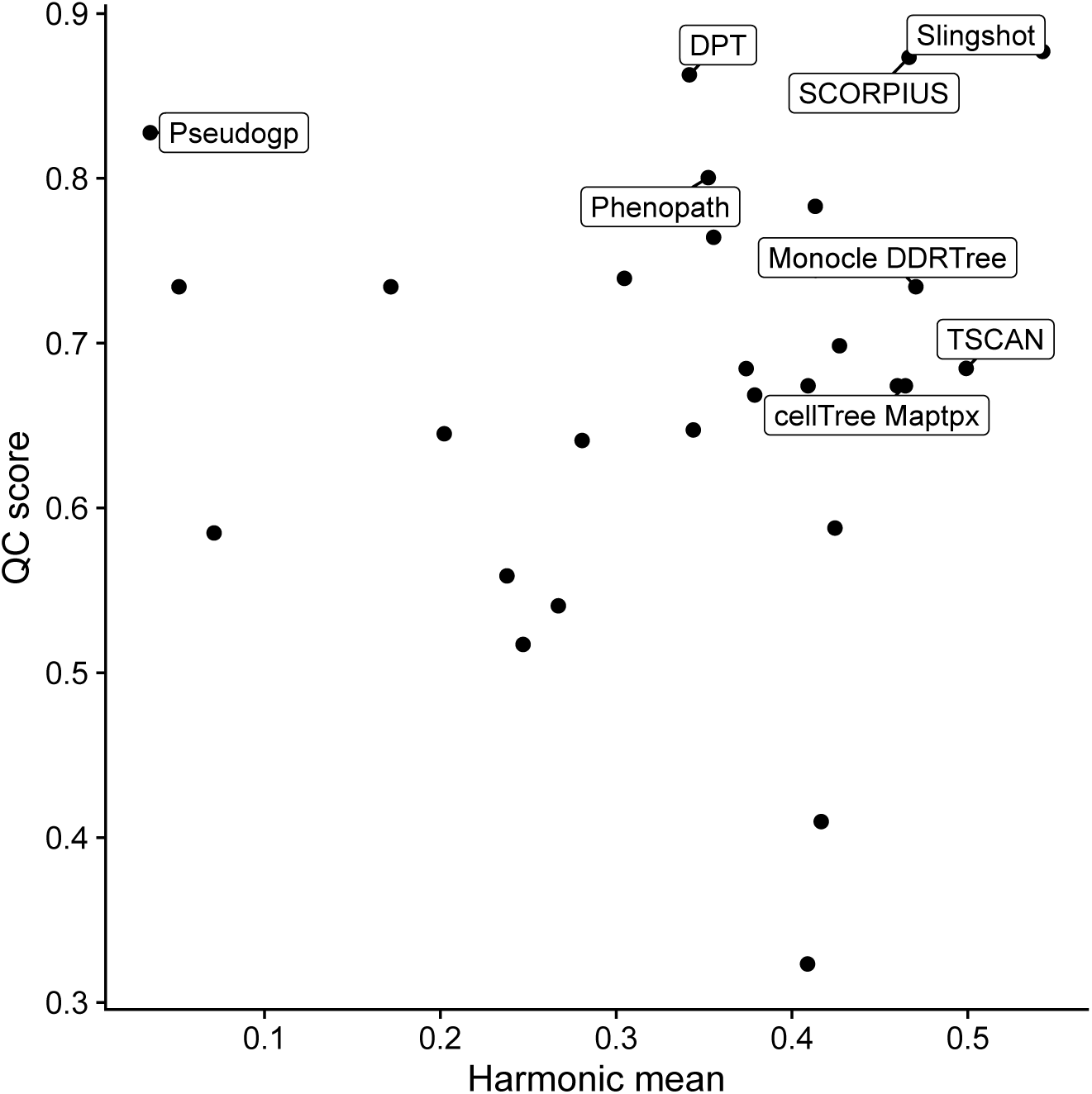
Comparison between method quality control scores and overall method performance.

**Supplementary Figure 15.**
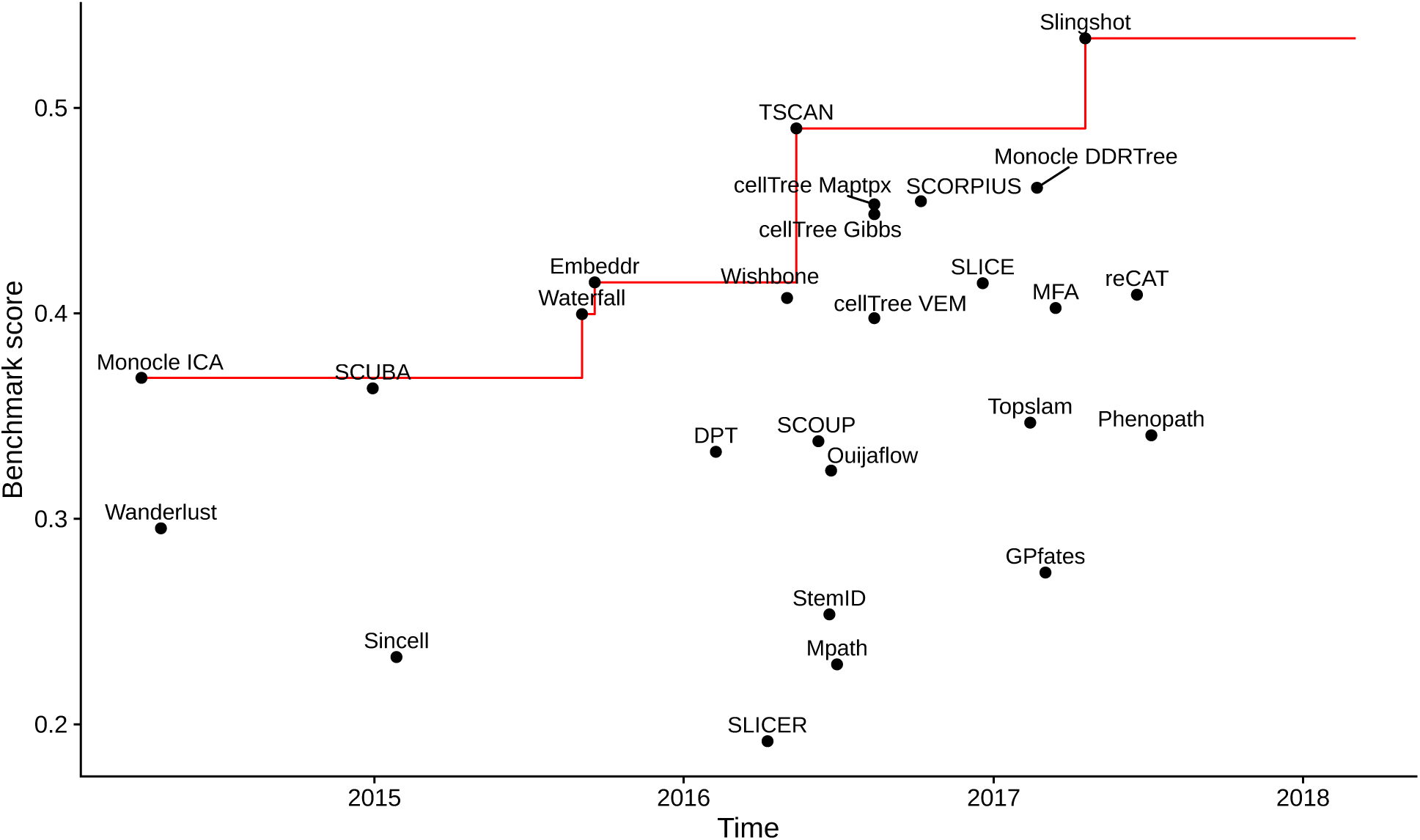
Method performance compared to the time the method was published. The red line indicates the optimal performance up to a point in time.

**Supplementary Figure 16.**
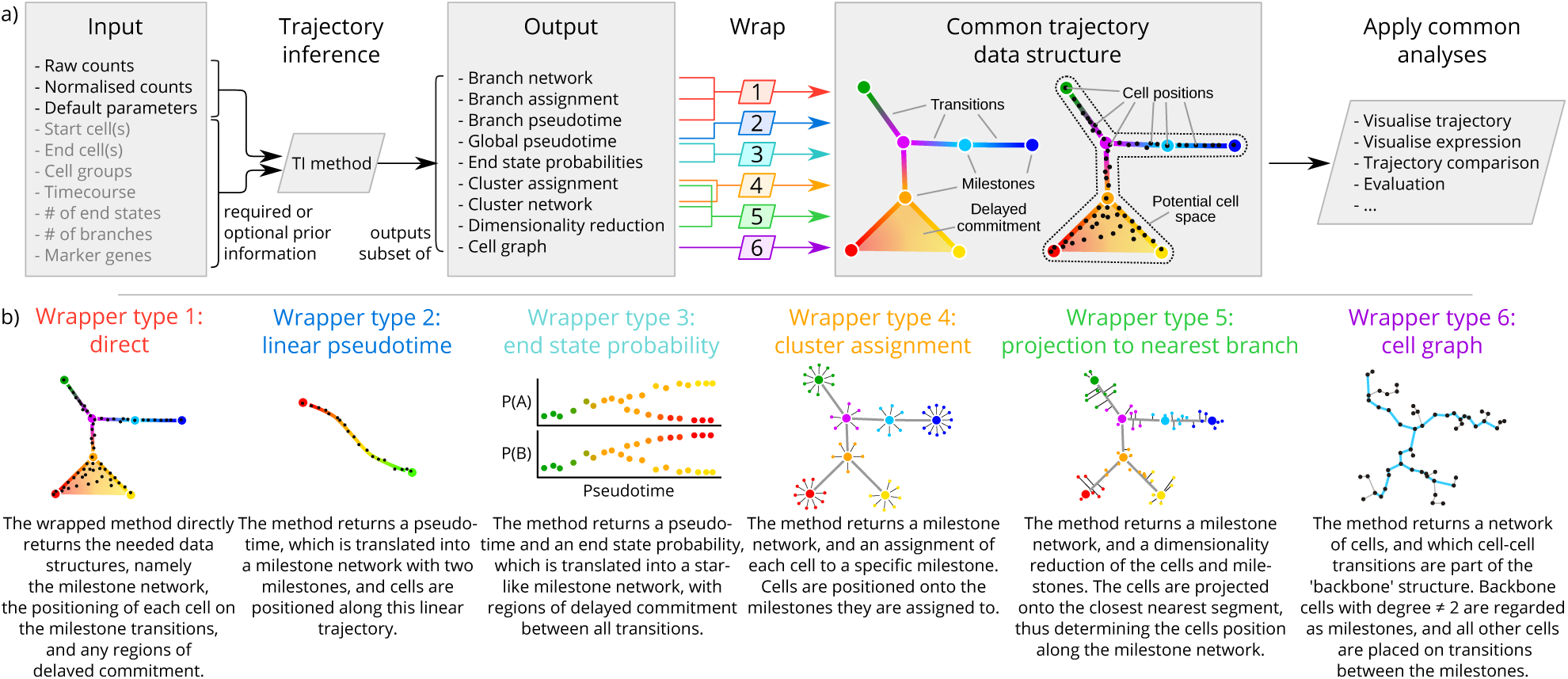
A common interface for TI methods. a) The input and output of each TI method is standardised. As input, each TI method receives either raw or normalised counts, several parameters, and a selection of prior information. After its execution, a method uses one of the six wrapper functions to transform its output to the common trajectory model. This common model then allows to perform common analysis functions on trajectory models produced by any TI method. b) The specific transformations performed by each of the wrapper functions is explained in more detail.

**Supplementary Figure 17.**
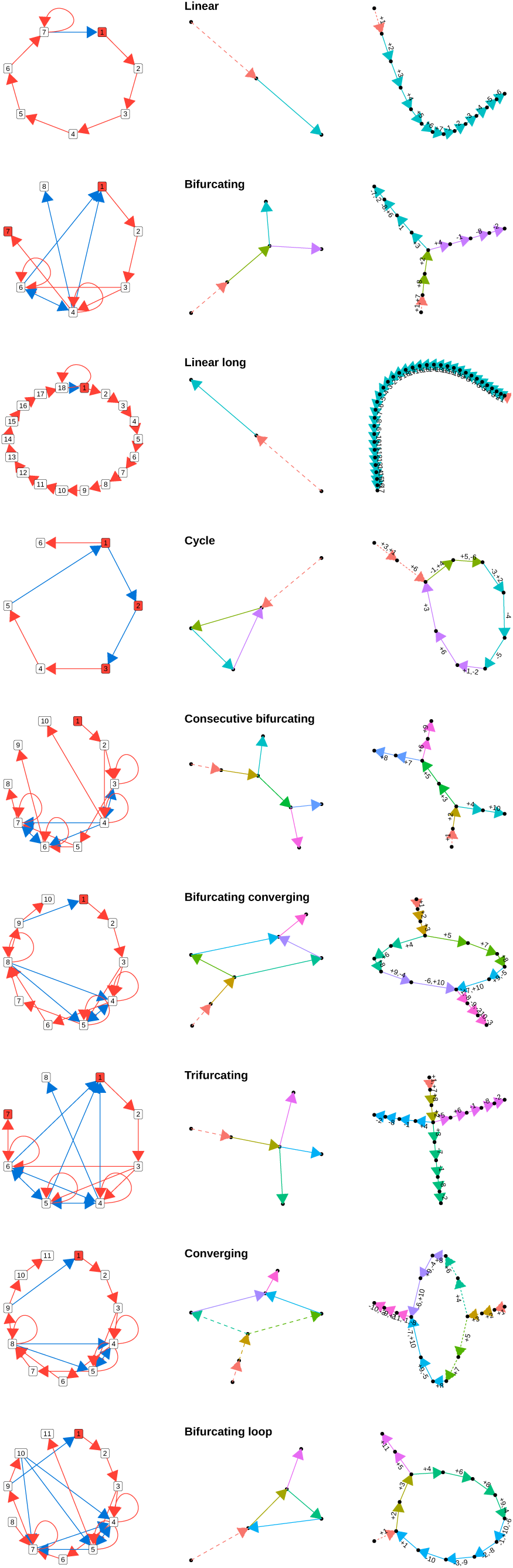
Module networks, milestone networks and the module dynamics for each of the different types of synthetic data

**Supplementary Figure 18.**
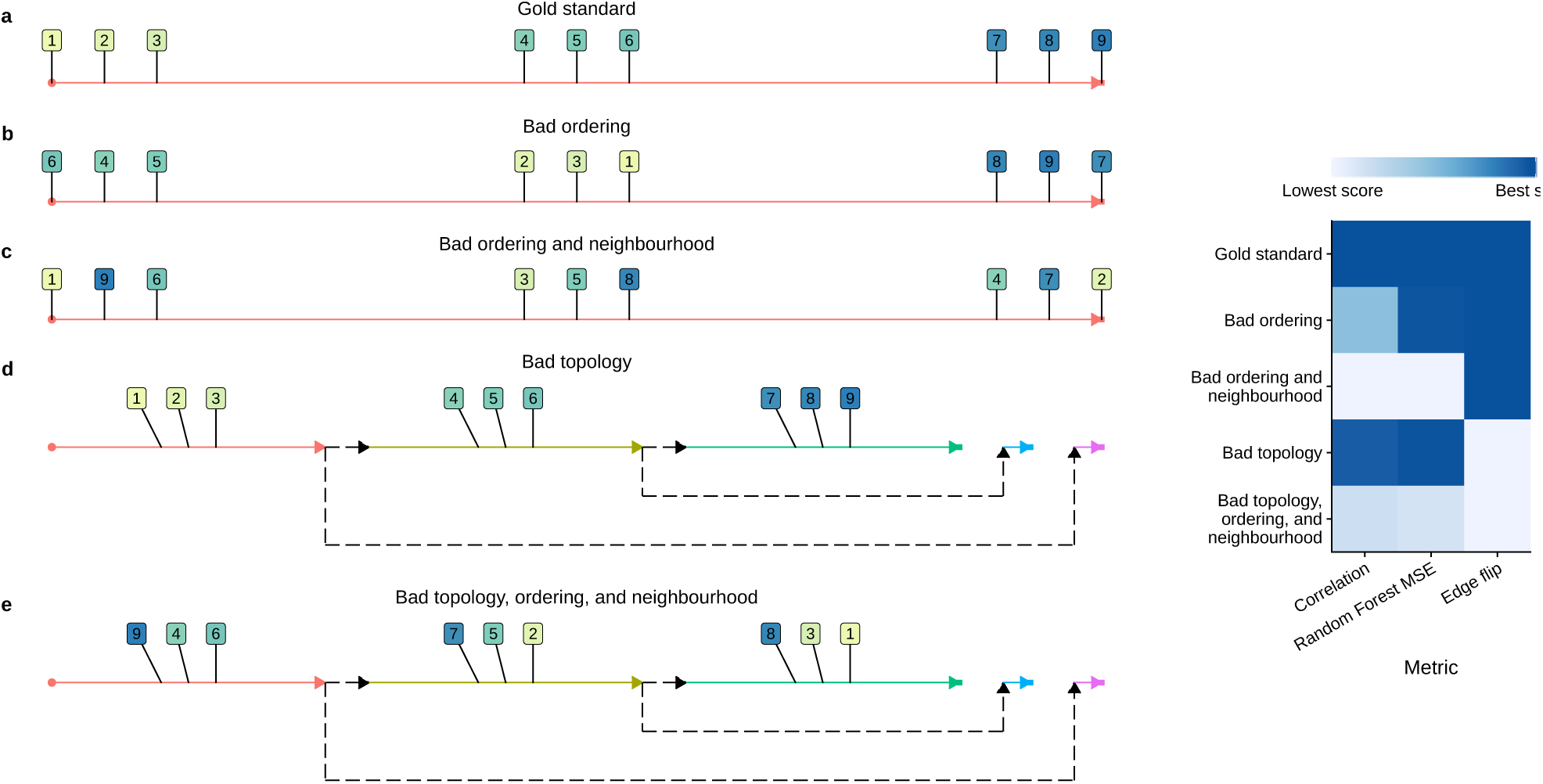
Different scores assess different aspects of the correspondence between a prediction and gold standard trajectory.

**Supplementary Figure 19.**
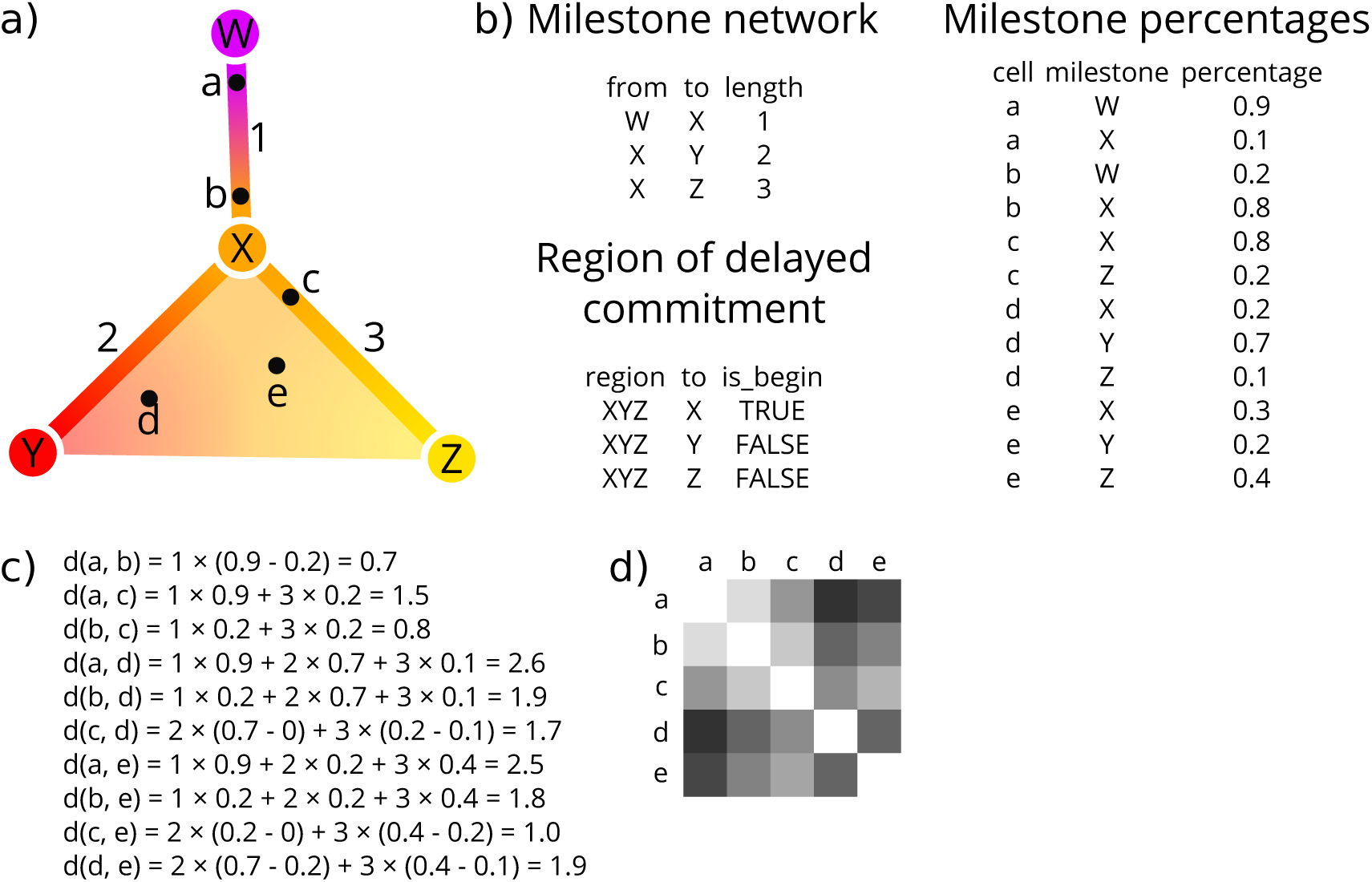
The modified geodesic distances demonstrated on a toy example. a) A toy example containing four milestones (W to Z) and five cells (a to e). b) The corresponding milestone network, milestone percentages and regions of delayed commitment, when the toy trajectory is converted to the common trajectory model. c) The calculations made for calculating the pairwise geodesic distances. d) A heatmap representation of the pairwise geodesic distances.

**Supplementary Figure 20.**
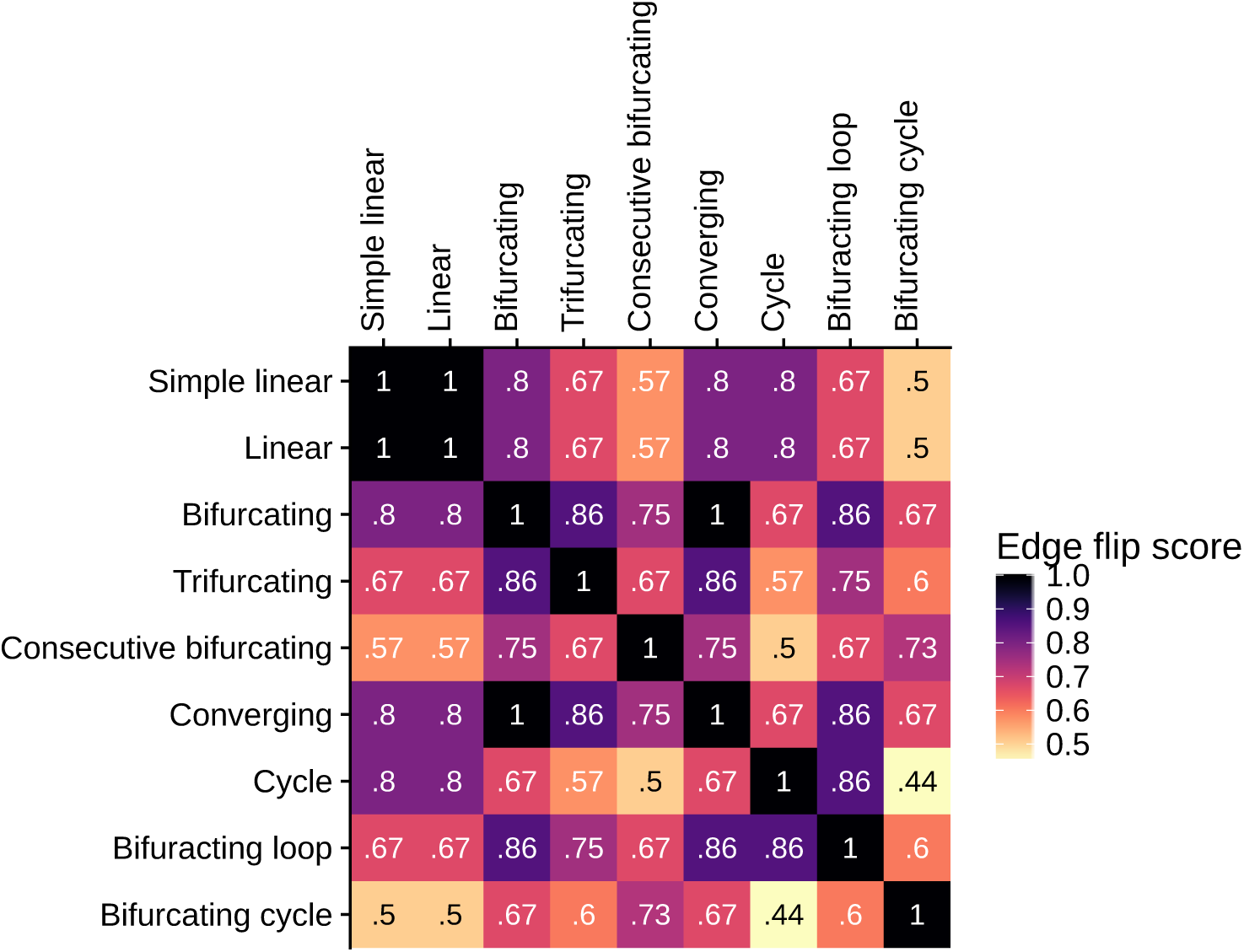
Edge flip score values for common trajectory topologies, defined in **Supplementary Figure 17**.

**Supplementary Figure 21.**
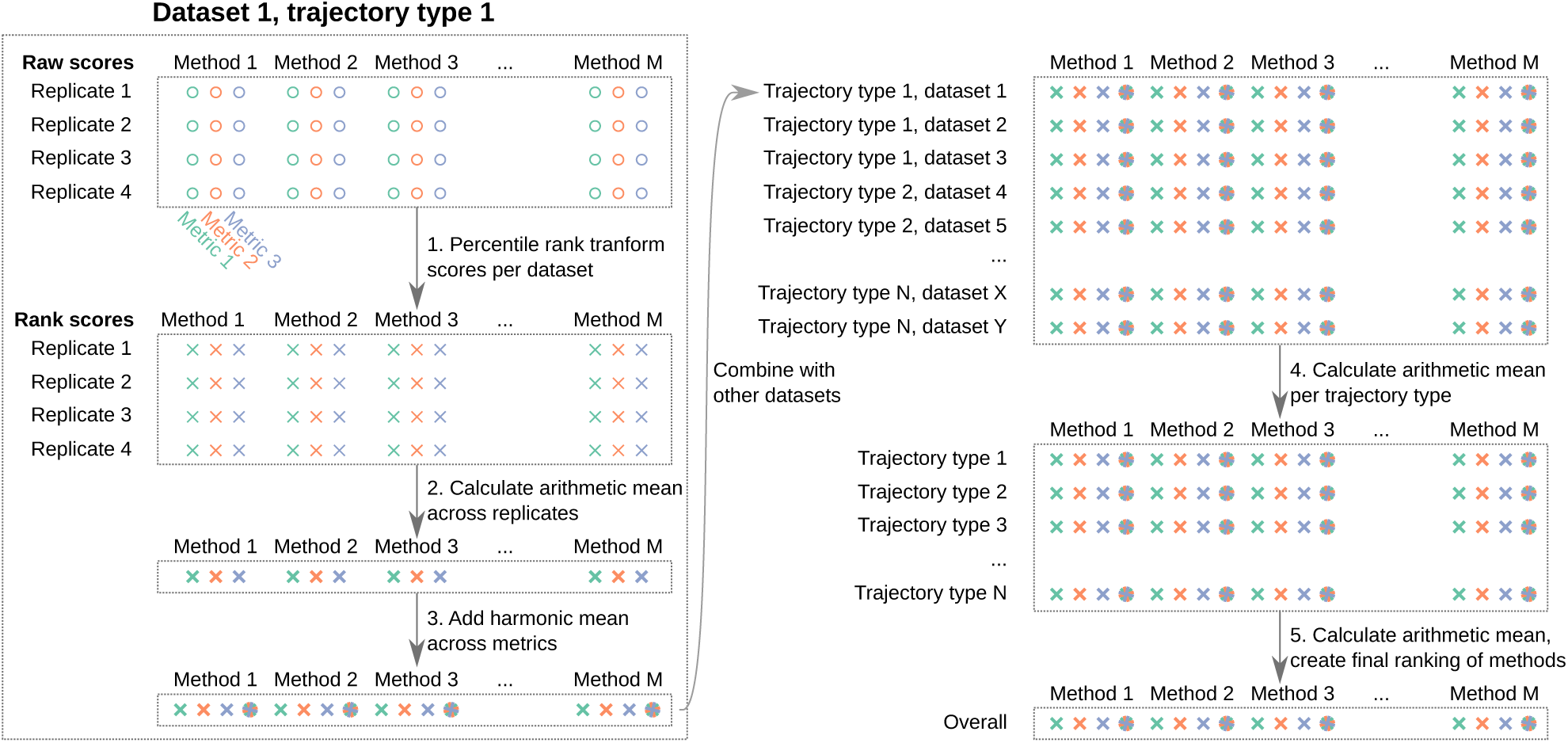
Aggregation methodology of metric scores.

**Supplementary Figure 22.**
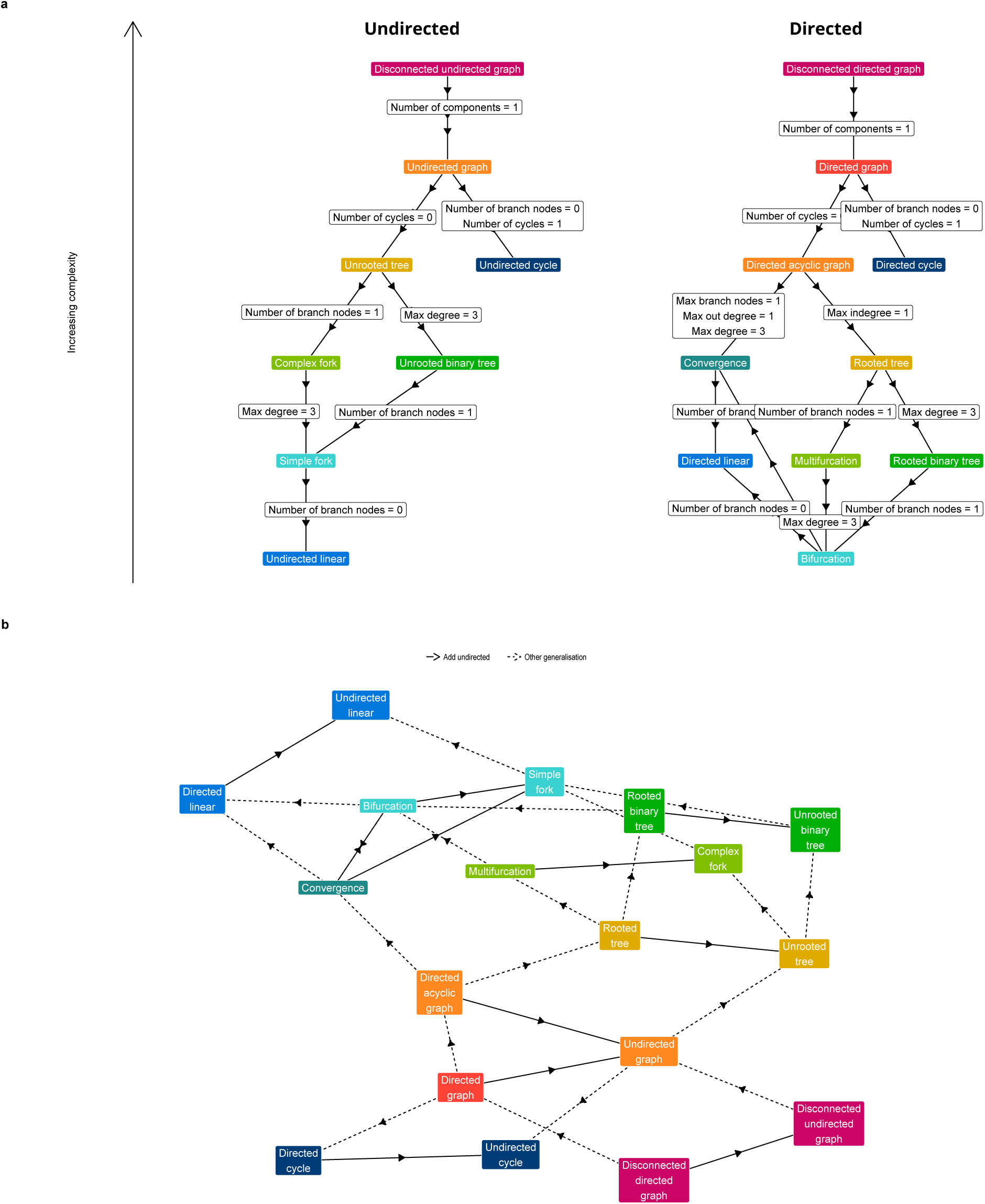
Different trajectory types can be classified in a directed acyclic graph structure, in which certain types are a generalisation of other trajectory types. a) The graph structure for undirected and directed trajectory types. Shown are the topological properties which change when going from a particular trajectory type to a less complex type. b) Combined structure for directed and undirected trajectory types.

## Supplementary Tables

**Supplementary Table 1.**
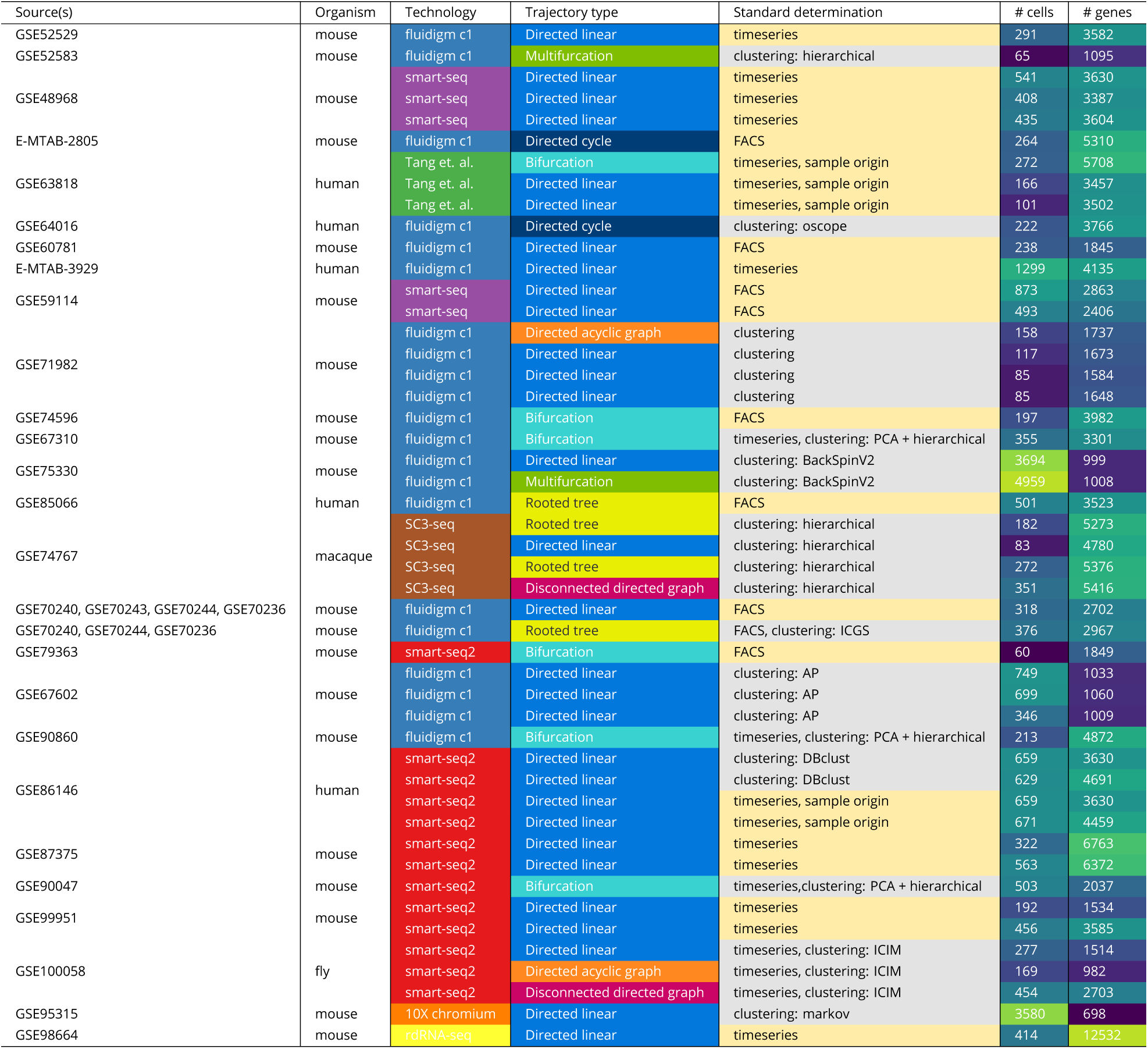
Real datasets used in this study.

**Supplementary Table 2.** Importance values for algorithmic components. The overall performance was compared between methods containing a particular algorithmic component, and we calculated both a corrected p-value using a two-tailed Mann– Whitney U test and the increase in node purity importance measure using random forest classification.

| Component | Adjusted p-value | Inc node purity | Effect | Methods | Category |
| --- | --- | --- | --- | --- | --- |
| principal curves | 0.08842 | 0.213 | Higher performance | SCUBA, Embeddr, SLICE, SCORPIUS, Slingshot | ordering |
| k-means | 0.08842 | 0.557 | Higher performance | Waterfall, SCORPIUS, Slingshot, Monocle DDRTree, reCAT | clustering |
| any graph building | 0.08842 | 0.739 | Higher performance | Monocle ICA, Waterfall, Wishbone, StemID, TSCAN, SLICER, Mpath, cellTree Maptpx, cellTree Gibbs, cellTree VEM, SLICE, Topslam, SCORPIUS, Slingshot, Monocle DDRTree, Sincell, Wanderlust, reCAT | graph building |
| MST | 0.13634 | 0.478 | Higher performance | Monocle ICA, Waterfall, StemID, TSCAN, Mpath, cellTree Maptpx, cellTree Gibbs, cellTree VEM, SLICE, Topslam, Slingshot, Monocle DDRTree, Sincell | graph building |
| tSNE | 0.13634 | 0.442 | Lower performance | StemID, Pseudogp, Topslam, Sincell | dimensionality reduction |
| any ordering | 0.13634 | 0.158 | Higher performance | Monocle ICA, SCUBA, StemID, TSCAN, SLICER, Embeddr, SLICE, SCORPIUS, Slingshot, Monocle DDRTree, Wanderlust | ordering |
| any clustering | 0.13634 | 0.113 | Higher performance | Waterfall, SCUBA, StemID, TSCAN, Mpath, SLICE, SCORPIUS, Slingshot, Monocle DDRTree, Sincell, reCAT | clustering |
| ICA | 0.31683 | 0.059 | Lower performance | Monocle ICA, Pseudogp, Topslam, Sincell | dimensionality reduction |
| any dimensionality reduction | 0.33783 | 0.251 | Higher performance | Monocle ICA, Waterfall, Wishbone, DPT, StemID, TSCAN, SLICER, Pseudogp, Embeddr, SLICE, Topslam, SCORPIUS, Slingshot, Monocle DDRTree, Sincell | dimensionality reduction |
| any dimensionality reduction | 0.33783 | 0.251 | Higher performance | Monocle ICA, Waterfall, Wishbone, DPT, StemID, TSCAN, SLICER, Pseudogp, Embeddr, SLICE, Topslam, SCORPIUS, Slingshot, Monocle DDRTree, Sincell | dimensionality reduction |
| knn graph | 0.39281 | 0.154 | Lower performance | Wishbone, SLICER, Sincell, Wanderlust | graph building |
| MDS | 0.61356 | 0.074 | Lower performance | Pseudogp, Topslam, SCORPIUS, Sincell | dimensionality reduction |
| any generative model | 0.73132 | 0.079 | Lower performance | GPFates, Pseudogp, SCOUP, cellTree Maptpx, cellTree Gibbs, cellTree VEM, MFA, Phenopath, SCIMITAR | generative model |
| PCA | 0.9791 | 0.033 | Lower performance | Waterfall, TSCAN, Pseudogp, SLICE, Topslam, Sincell | dimensionality reduction |
| k-medoids | 0.9791 | 0.057 | Lower performance | StemID, SLICE, Sincell, reCAT | clustering |

**Supplementary Table 3.**
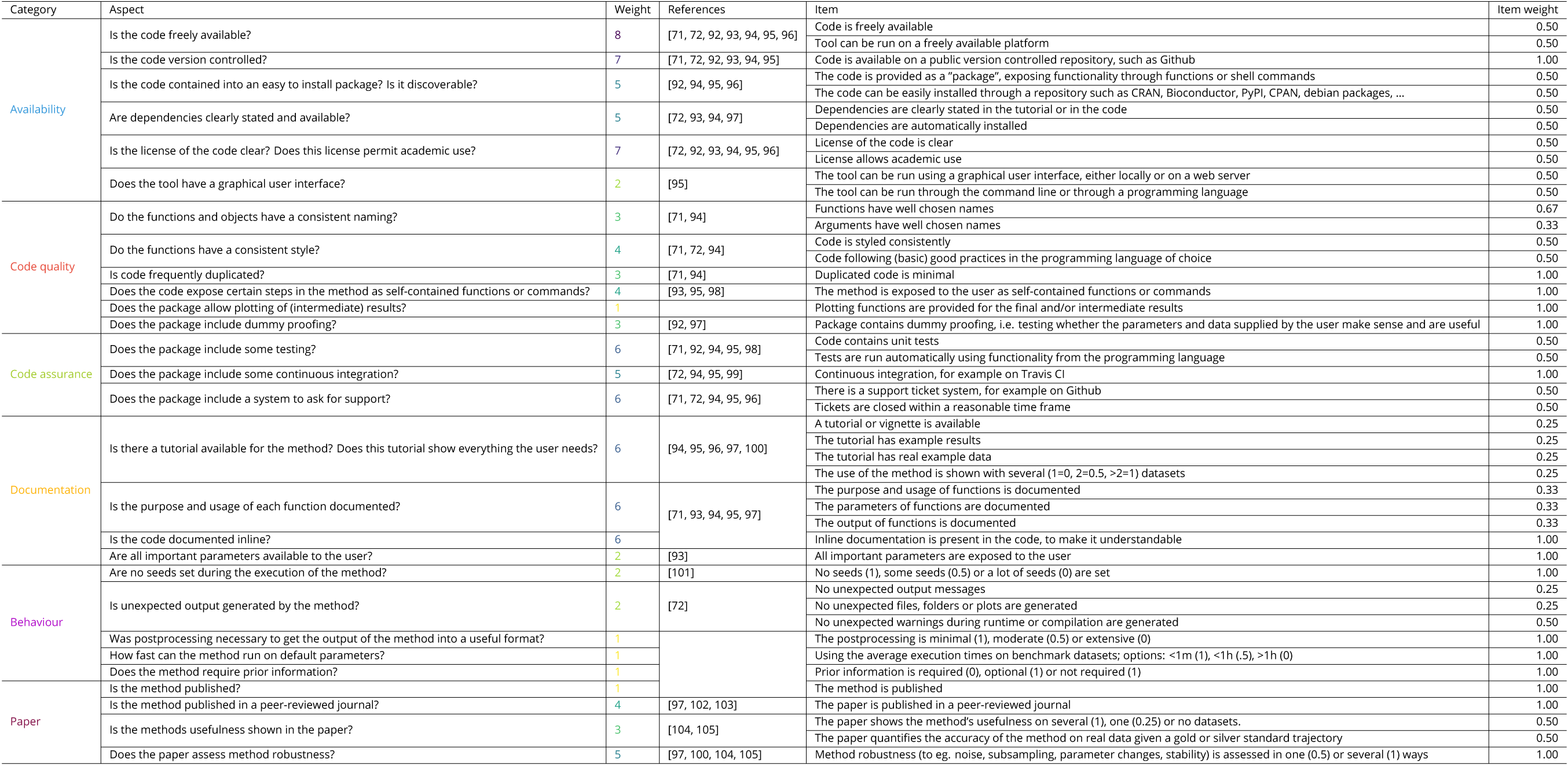
Scoring scheme for method quality control. Each quality aspect was given a weight based on how many times it was mentioned in a set of articles discussing best practices for tool development.

**Supplementary Table 4.** Distributions from which each parameter in the thermodynamic model was sampled.

|  |
| --- |
| $r = \mathcal{U}(10, 200)$ |
| $d = \mathcal{U}(2, 8)$ |
| $p = \mathcal{U}(2, 8)$ |
| $q = \mathcal{U}(1, 5)$ |
| $a_0 = \begin{cases} 1 & \text{if } e \in \{1\} \text{ or } e = \emptyset \\ 0 & \text{if } e \in \{-1\} \\ 0.5 & \text{otherwise} \end{cases}$ |
| $a_i = \begin{cases} 1 & \text{if } e_i \in \{-1\} \\ 0 & \text{otherwise} \end{cases}$ |
| $s = \mathcal{U}(1, 20)$ |
| $k = y_{max}/(2 * s)$ |
| $c = \mathcal{U}(1, 4)$ |
| **where** |
| $y_{max} = r/d * p/q$ |

## Colophon

This report was generated on 2018-03-05 23:36:32 using R version 3.4.3 (2017-11-30) and the following packages:

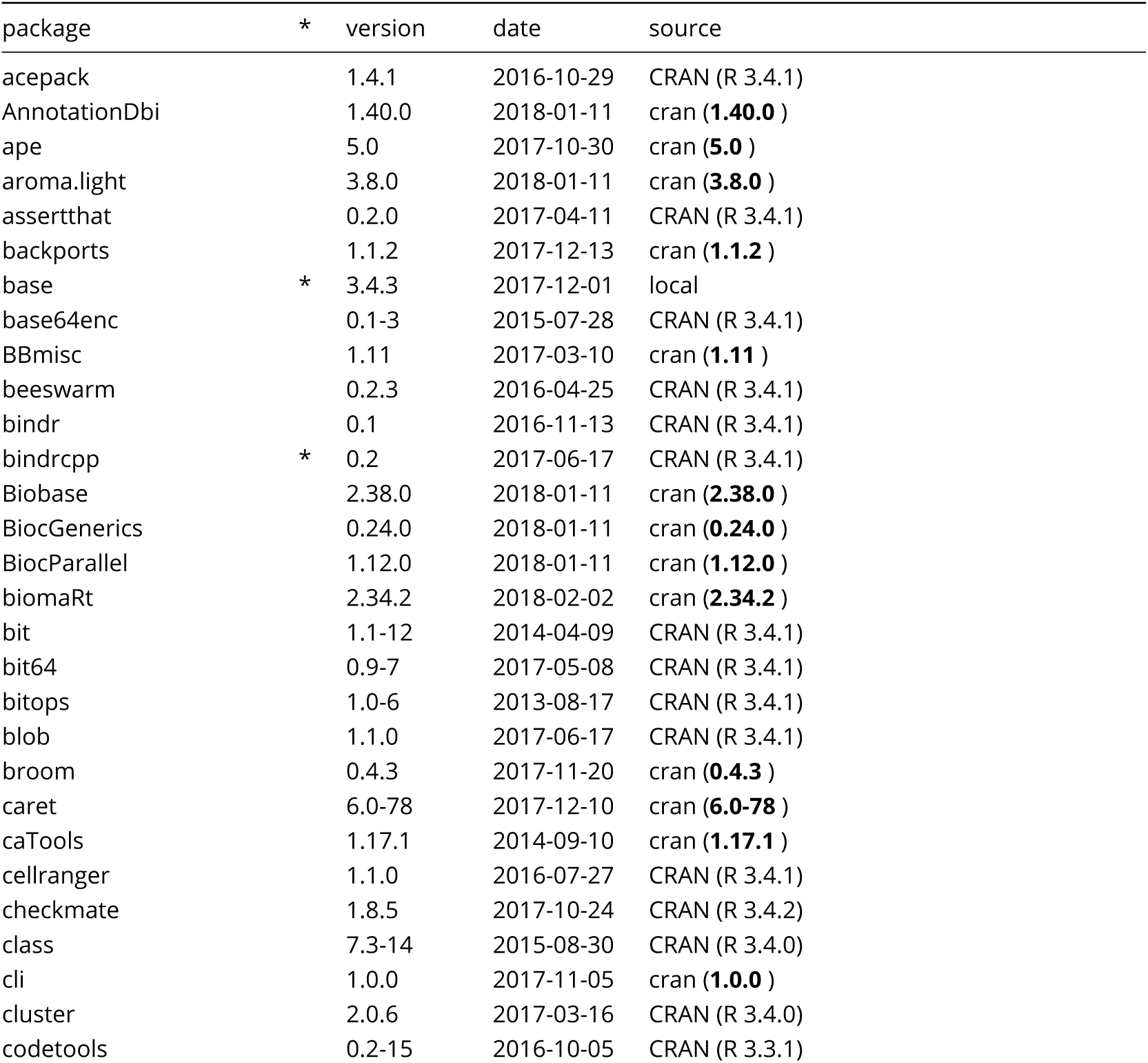
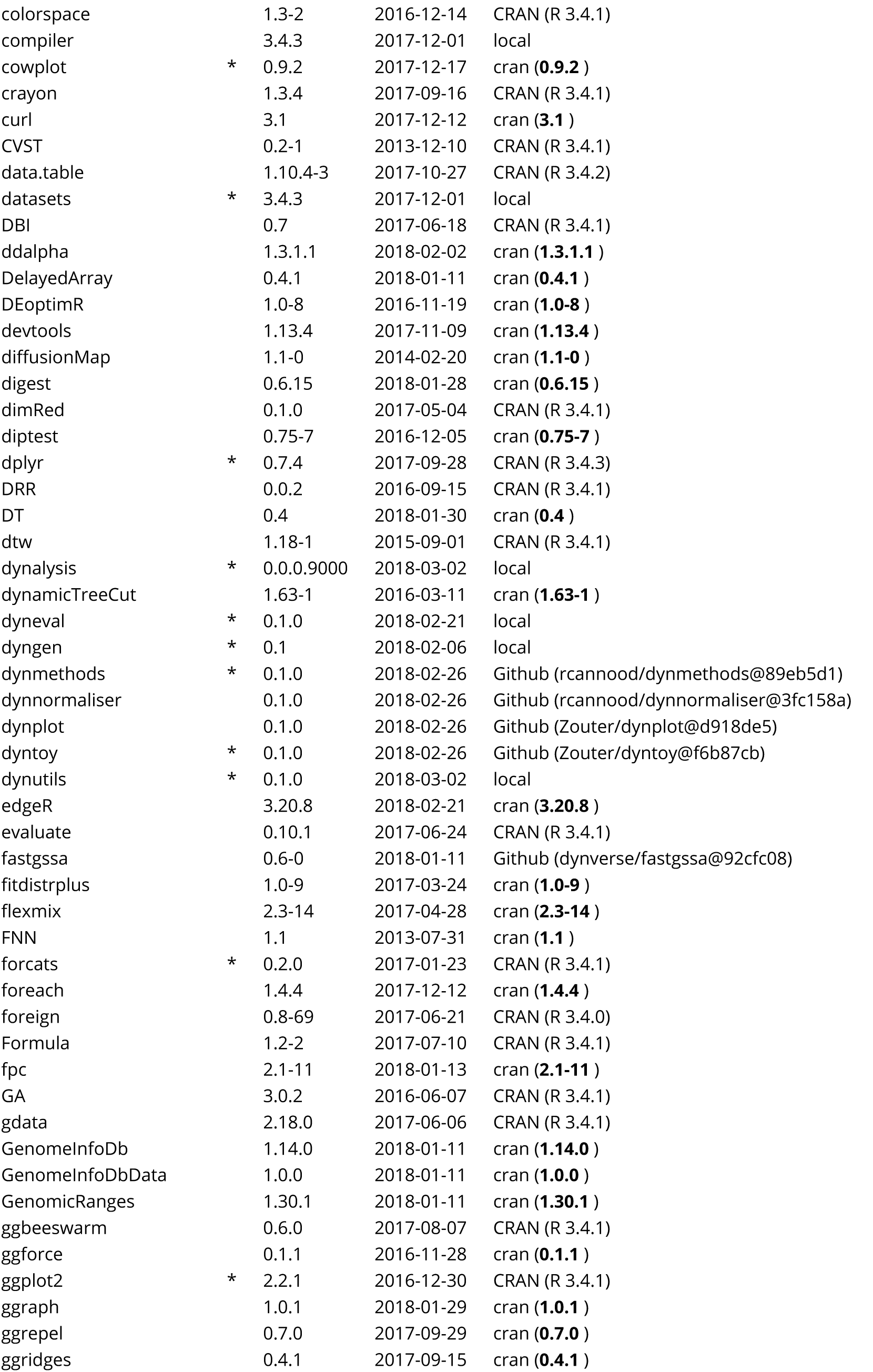
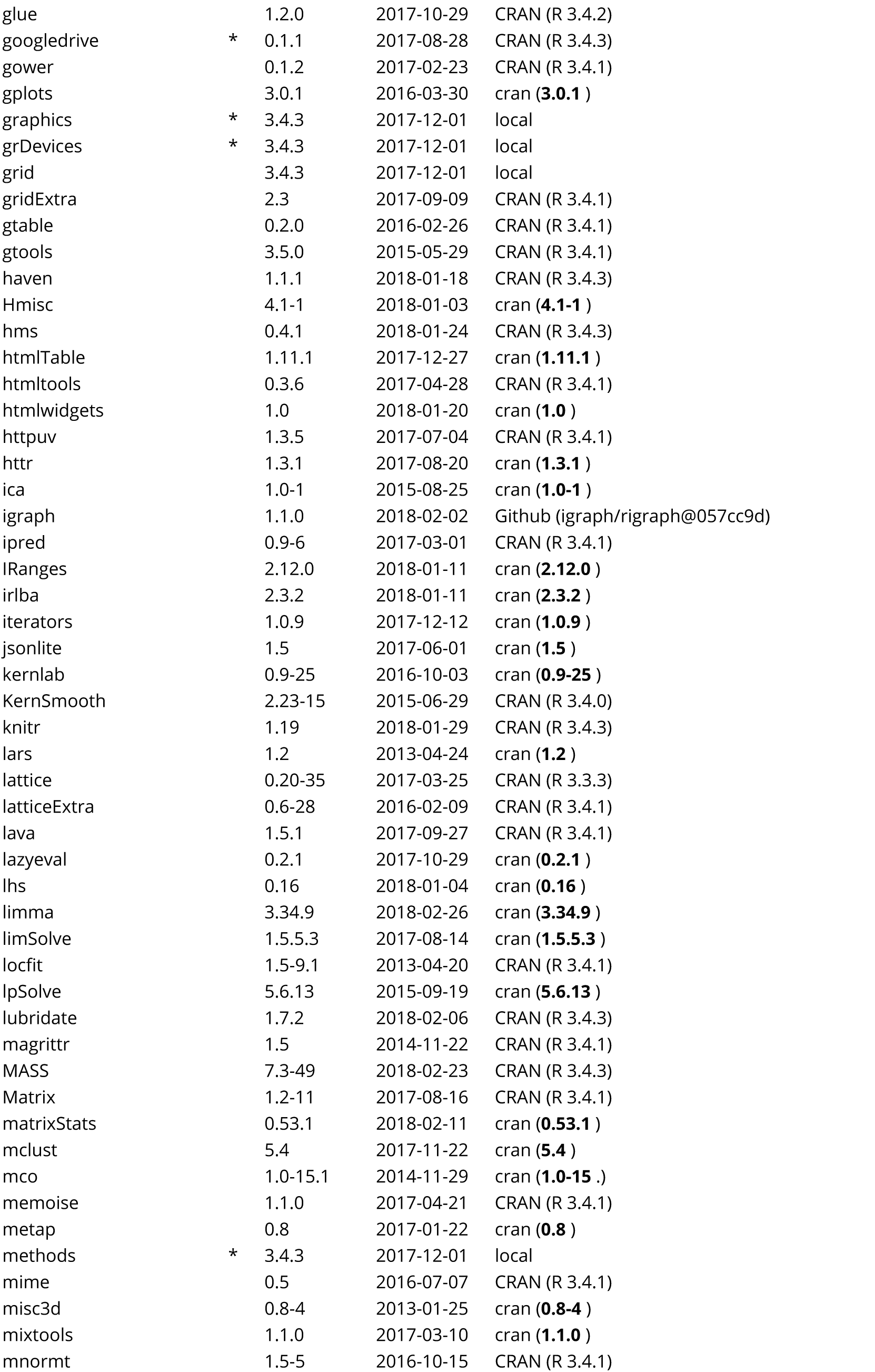
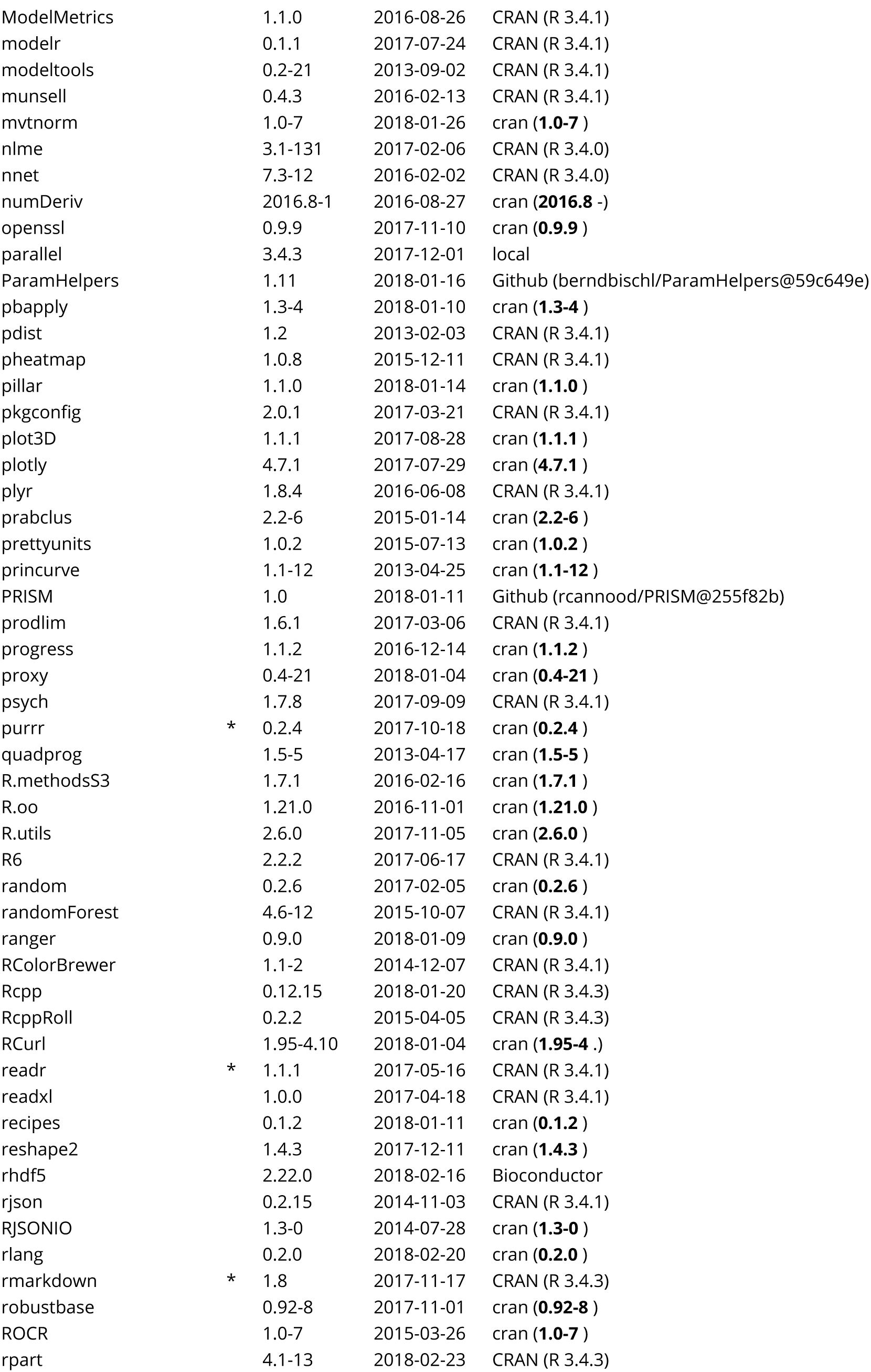
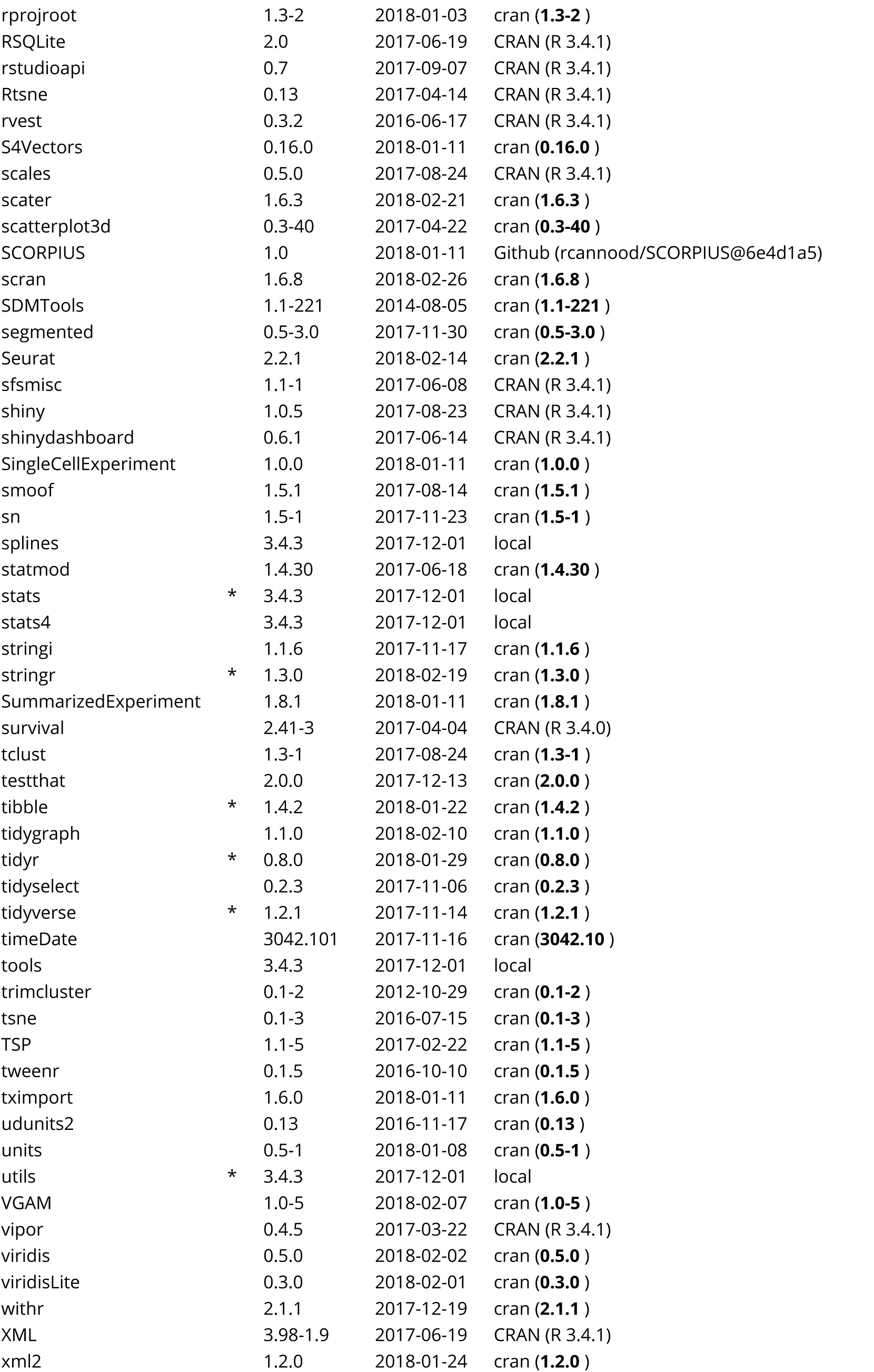
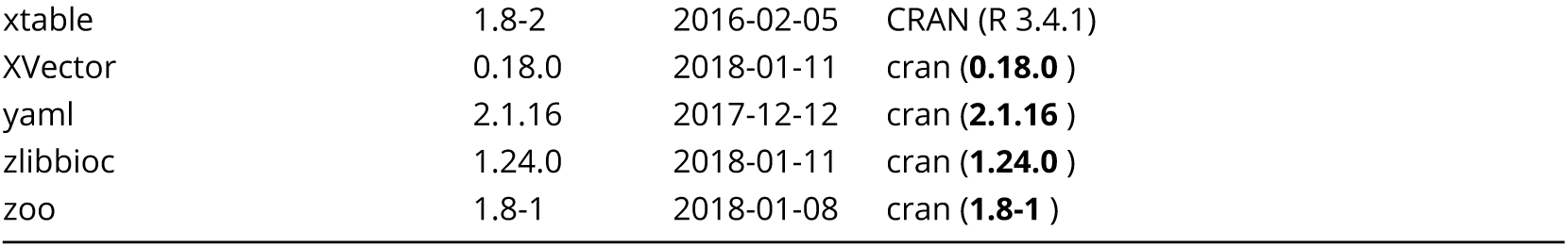

